# In-cell architecture of the mitochondrial respiratory chain

**DOI:** 10.1101/2024.09.03.610704

**Authors:** Florent Waltz, Ricardo D. Righetto, Ron Kelley, Xianjun Zhang, Martin Obr, Sagar Khavnekar, Abhay Kotecha, Benjamin D. Engel

## Abstract

Mitochondria produce energy through oxidative phosphorylation, carried out by five membrane-bound complexes collectively known as the respiratory chain. These complexes work in concert to transfer electrons and pump protons, leading to ATP regeneration. The precise organization of these complexes in native cells is debated, notably their assembly into higher-order supercomplexes called respirasomes. Here, we use *in situ* cryo-electron tomography to visualize the native structures and organization of several major mitochondrial complexes inside *Chlamydomonas reinhardtii* cells. ATP synthases and respiratory complexes are segregated into curved and flat crista membrane domains, respectively. Respiratory complexes I, III, and IV assemble into a single type of respirasome, from which we determined a native 5 Å-resolution structure showing the binding of electron carrier cytochrome *c*. Combined with single-particle cryo-electron microscopy reconstruction at 2.4 Å resolution, we assemble a detailed model of how the respiratory complexes interact with each other inside native mitochondria.

## Introduction

Mitochondria are essential organelles found in nearly all eukaryotes (*1*). One of their main functions is oxidative phosphorylation, where a set of five membrane-embedded molecular machines, the respiratory complexes, work together to recycle ADP to ATP, the energy currency of the cell (*2, 3*). Even though many high-resolution structures of isolated respiratory complexes have been solved (*4–10*), their organization within mitochondria remains an ongoing matter of investigation. They localize to cristae, specialized inner membrane (IM) folds that greatly increase the membrane surface area (*11*). The complexes can form higher-order structures, including respirasome supercomplexes (containing complexes I, III, IV, and sometimes II) (*12–22*) and oligomers of ATP synthases (*11, 23*), both of which can influence crista membrane architecture (*11*). However, respirasomes have only been described from biochemically purified complexes or membranes, using techniques such as blue-native PAGE (*24–27*) or single particle analysis (SPA) of cryo-electron microscopy (cryo-EM) data (*12–22*). Purification of membrane complexes typically involves detergents, which can extract structural lipids and cause rearrangements. Moreover, the disruption of the membranes during isolation of mitochondria might have a direct impact on respiratory chain function, e.g., releasing the electron carrier cytochrome *c* (cyt. *c*) that shuttles within the intermembrane space (IMS) between complexes III and IV.

To better understand the molecular organization and function of respirasomes, these supercomplexes should be studied *in situ* within their native cellular environment. We therefore sought to visualize native mitochondria using cryo-electron tomography (cryo-ET) (*28, 29*), which recently has experienced breakthroughs in resolution thanks to rapid development of both hardware and software (*30–32*). With this approach, frozen cellular samples are thinned using cryo-focused ion beam milling (cryo-FIB) to produce 100-200 nm-thick sections, which are then imaged in 3D using a transmission electron microscope (*33, 34*). We chose to work with *Chlamydomonas reinhardtii*, a unicellular green alga widely used to study photosynthesis (*35*) and ciliary function (*36, 37*) which has also proven to be an excellent model to investigate mitochondrial biology (*38, 39*). High-throughput milling with a cryo-plasma FIB enabled us to acquire a large scale cryo-ET dataset of *C. reinhardtii* cells (EMPIAR-11830). This enabled us to visualize the native structures and molecular organization of several mitochondrial complexes, including a 5 Å-resolution structure of a respirasome within the native cellular environment. Compared to a structure of an isolated partial respirasome we generated using SPA, our *in situ* structure reveals key conformational differences and interaction interfaces, as well as the binding sites of cyt. *c*, demonstrating the power of in-cell structure determination.

## Results & Discussion

### Molecular landscape of mitochondria revealed by cryo-ET

In *C. reinhardtii,* mitochondria form an almost uninterrupted network that covers the entire cell and can be reorganized upon different conditions (*40*). Cryo-ET of mitochondria inside *C. reinhardtii* cells allowed direct visualization of their native membrane architecture and molecular organization. A prominent architectural feature of mitochondria are the cristae. Like in animals, the cristae in Chlamydomonas are shaped like flat lamellar sacs. The cristae lumen in *C. reinhardtii* has an average width of ∼15 nm, compared to ∼12 nm measured in several animal studies (*41–43*). Visual inspection of the tomograms revealed several mitochondrial complexes of known and unknown identity, which we then structurally resolved by subtomogram averaging (STA) (Fig. 1 and fig. S1).

**Figure 1:**
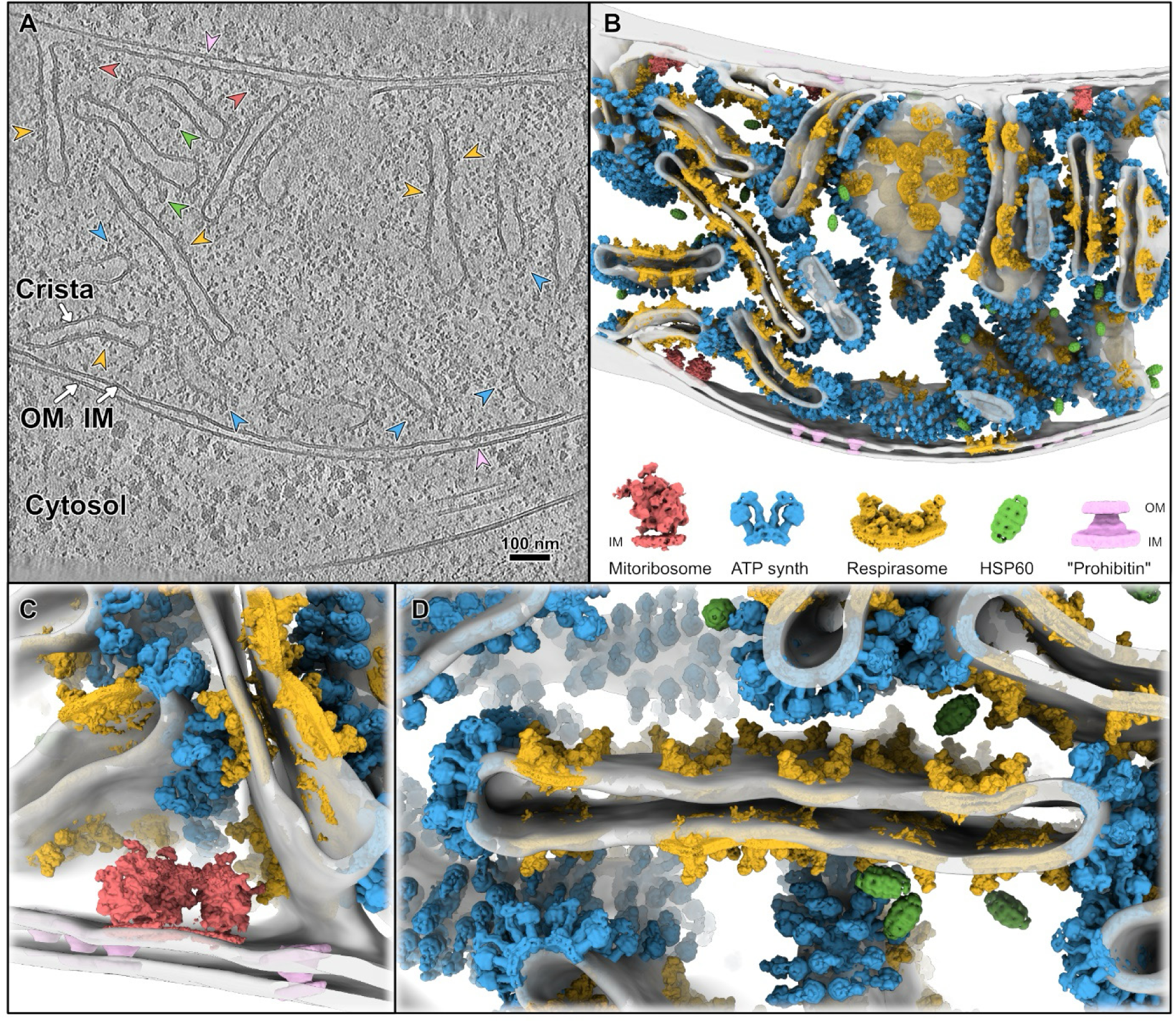
Molecular landscape of Chlamydomonas mitochondria. **A)** Slice through a tomogram depicting a mitochondrion within a native *C. reinhardtii* cell. **B)** Corresponding segmentation of the tomogram with STA structures of the molecular complexes mapped back into the respective particle positions. Grey: mitochondrial membranes, red: mitoribosomes, blue: ATP synthase dimers, orange: respirasomes, green: HSP60, pink: putative prohibitins. Enlarged views of the STA maps are displayed below the segmentation. **C and D)** Close-up views of the mapped in complexes, showing the organization of mitoribosomes and putative prohibitins at the inner membrane (C) and the organization of ATP synthases and respirasomes on cristae (D).

At the IM, we observed that mitochondrial ribosomes were often organized as circular polysomes, a feature that was not previously described (fig. S1B) (*38*). Within the heavily crowded mitochondrial matrix, we identified “bullet-shaped” complexes with 7-fold symmetry that could be attributed to HSP60 after STA. In the IMS, between the IM and outer membrane (OM), we observed multiple “volcano-shaped” complexes (Fig. 1B and fig. S1A). These complexes are excluded from cristae and maintain a consistent orientation, with the wide edge touching the IM and the narrow tip connecting to the OM. The identity of these complexes is uncertain, but their structure is similar to prohibitin multimers, ubiquitous among eukaryotes including Chlamydomonas, and recently described in human cells (*44*). A rarer observation (about 1 per tomogram) were “lightbulb-shaped” complexes bound to the OM and protruding into the cytosol (fig. S1D). We were not able to clearly average these structures due to the limited number of particles and probable structural heterogeneity of the complexes.

We then analyzed the organization of the complexes located on the crista membranes. At the tips of the cristae, ATP synthases could be clearly identified (Fig. 1D), forming dimers that organize into rows (fig. S1E). We observe that these ATP synthase dimer rows form higher-order structures, with multiple rows aligned next to each other, thereby inducing the complete fold of the crista tip (fig. S1E). This organization agrees with the arrangement of ATP synthase recently reported in the non-photosynthetic green alga Polytomella (*45*). This is however a distinct cristae-shaping mechanism compared to yeast and animals, where a single row of ATP synthase dimers is supposedly enough to induce the full fold of the cristae membrane (*46, 47*), and Toxoplasma (*5*), where hexameric ATP synthases shape the tips of tubular cristae.

### Subtomogram average of the native Chlamydomonas respirasome

Densities were also visible on flat regions of crista membranes, where respiratory complexes involved in proton translocation are supposed to be located. Indeed, complex I could be clearly identified by eye (Fig. 1A). We used template matching to detect either complex I or complex III (see Methods). After extraction and classification of the particles, both complex I and III template matching results converged towards a similar class, a respirasome supercomplex harboring a C2 symmetry (fig. S2). This low-resolution average of the supercomplex was then used as a reference to template match the entire set of mitochondria-containing tomograms. Subtomogram classification and refinement rounds (fig. S3) with and without symmetry imposition, allowed us to reach a global resolution of 5.4 Å from 14,494 particles. The membrane-embedded part of complex I was best-resolved (local resolution reaching 4.1 Å), indicating that the transmembrane domains of complex I make stable interactions within the supercomplex. To further improve the map, we recentered the alignment on one asymmetric half of the respirasome and used symmetry expansion. This yielded a 6.2 Å map, which while slightly lower resolution than with C2 symmetry applied, was more homogeneously resolved throughout all the protein complexes. Inspection of the respirasome structure allowed us to determine that it is composed of two monomers of complex I, two dimers of complex III, and six monomers of complex IV. This I_2_ III_4_ IV_6_ respirasome (Fig. 2) has a distinct arrangement of respiratory complexes compared to supercomplexes previously described from other organisms. The local resolution of the different complexes (Fig. 2B) allowed us to observe bulky side chains in the transmembrane core of complex I, and even to determine that the catalytic site of the carbonic anhydrase domain is active in Chlamydomonas. We could also resolve bound heme groups in both complexes III and IV.

**Figure 2:**
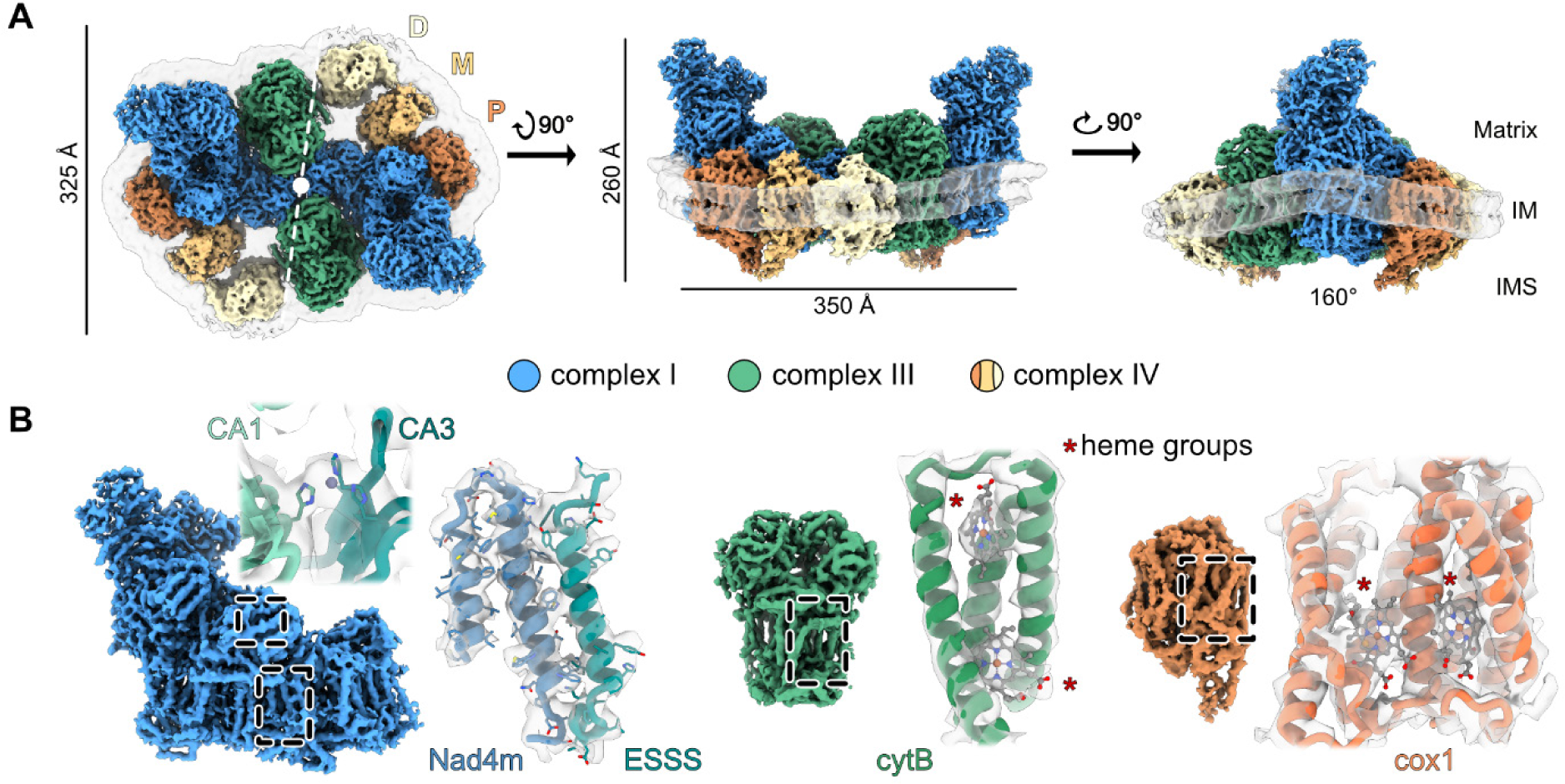
Subtomogram average of the native Chlamydomonas respirasome. **A)** STA maps of the *C. reinhardtii* mitochondrial respirasome. From left to right: view from the matrix side (white dashed line: 2-fold symmetry axis) and two different side views. Bending of the IM (transparent grey) is shown in the rightmost side view. Blue: complex I, green: complex III, brick/yellow/cream: complex IV. P, M, and D indicate proximal, middle, and distal complex IV monomers. **B)** Close-up views of the different features resolved in the STA map of each complex. Left: active site of the carbonic anhydrase and bulky side chains in the membrane arm of complex I; Middle and right: heme groups in complexes III and IV.

The I_2_ III_4_ IV_6_ respirasome is ∼260 Å in height, ∼335 Å in width, and ∼345 Å in length, and contains 384 transmembrane helices (Fig. 2A and Movie 2). The 2-fold symmetry axis is positioned at the interface between the two monomers of complex I, mediated by the tips of the distal membrane domains. On both sides of complex I, each complex III dimer interacts with the matrix-exposed distal membrane domain, locking the two complex I monomers together. We have designated the complex IV monomers as proximal (P), middle (M), and distal (D) for their positions relative to complex I (Fig. 2A). The two best resolved complex IV monomers (P and M) are anchored directly next to complex I. The most proximal complex IV (P), closest to the carbonic anhydrase domain of complex I, is oriented with its concave side facing complex I. The middle complex IV (M) is rotated by about 90°, and has its concave side facing the proximal complex IV, while still interacting with complex I. The distal complex IV (D) is oriented in a similar fashion as the middle complex IV; however, it is not directly interacting with complex I but appears to form some contacts with complex III from its convex side. This complex IV exhibited a fainter density in the STA map compared to the other two, which may suggest that Chlamydomonas respirasomes are not always bound to the distal complex IV *in vivo*. An alternative explanation for the fainter density could be structural flexibility, as the distal complex IV makes limited contacts with the other complexes of the respirasome, which suggests a looser interaction and hence conformational heterogeneity. The respirasome induces local membrane bending with an angle of ∼160° (Fig. 2A), similar to the bending angle formed by CI + CIII_2_ in Arabidopsis (*18, 20*). However, the bending induced by a single respirasome does not appear to have an impact on overall curvature of the surrounding crista membrane, which is flat.

Although the majority of respirasomes were localized to the flat crista membranes, a subpopulation was also found at the IM, excluded from cristae (see Fig. 1B and fig. S2C). Subunits of complexes I and IV are encoded in the mitochondrial genome and are first synthesized at the inner membrane, where mitochondrial translation takes place (*38, 48, 49*). Thus, these supercomplexes might be newly synthetized ones, on their way to relocate to the cristae. Alternatively, they might contribute to a local electrical potential gradient, *Δψ*, necessary for ATP-independent protein import machinery at the IM (*50*). This observation suggests that respiratory complexes may function not only in cristae but also at the inner membrane.

This is, to date, the only structure of respirasomes inside native cells. Previous *in situ* cryo-EM and cryo-ET of respirasomes in mammals (*51*), Tetrahymena (*17*), yeast, and plants (*52*) were performed on isolated membranes or mitochondria. Here, we resolve one main respiratory supercomplex, with distinct stoichiometry from the multiple types of respirasomes recently observed in purified pig heart and mouse mitochondria (*12, 51*). The Chlamydomonas cells we imaged were all grown asynchronously under similar conditions, so we cannot exclude that different respirasomes might form in other conditions to meet changing metabolic needs. It might be that the supercomplex we observed has the optimal stoichiometry or arrangement for a specific metabolic state. Individual complex I and complex III were not detected by our template matching approach, and thus may not be abundant *in situ*. Even if the free states do exist, we expect complexes I, III, and IV to directly assemble into supercomplexes, probably during the biogenesis of each complex. However, our dataset is not well suited to study these events, as we primarily imaged mature mitochondria where the synthesis of new complexes occurs infrequently.

### Integration with single-particle cryo-EM

To better interpret the composition and inter-subunit interactions of our in-cell structure of the Chlamydomonas respirasome, we also biochemically isolated a supercomplex from *C. reinhardtii* and determined its structure at 2.8 Å resolution using single-particle cryo-EM. The isolated supercomplex is composed of one complex I, one dimer of complex III, and two monomers of complex IV (I_1_ III_2_ IV_2_ stoichiometry). Except for one missing complex IV (from the D position), it is almost identical in composition and conformation to half of a native respirasome, consistent with the pseudo-C2 symmetry observed *in situ*. Local refinements reached an average of 2.3 Å on complex I, 2.4 Å on complex III, and 2.4 Å on complex IV (fig. S4). This allowed us to sequence the proteins directly from the SPA map and a build a full model *de novo* (Fig. 3A). In total, Chlamydomonas complex I is composed of 51 proteins, complex III of 20 proteins (10 unique, considering that complex III is a homodimer) and complex IV of 12 proteins (see figs. S5-S7 and Tables 2 and 3 for a complete description of the proteins). Based on their protein composition, we determined the molecular weight of these complexes, with complex I at 1640 kDa, complex III at 461 kDa and 198 kDa for complex IV, which makes the isolated I_1_ III_2_ IV_2_ a total of ∼2.5 MDa with 95 protein subunits, and the native I_2_ III_4_ IV_6_ a total of ∼5.4 MDa with 214 protein subunits. Briefly, complex I structure and composition is similar to what was previously described in flowering plants and non-photosynthetic algae (*10*), but here we identify all the different protein subunits, including protein P10 at the very tip of the distal membrane arm, which appears to be a frame-shifted version of the predicted Chlre_08g368050v5 open reading frame. Interestingly, the carbonic anhydrase domain, originally described in plants, and now established as an ancestral domain of mitochondrial complex I (*53*), is active, as attested by the presence of a zinc ion at the interface of the CAG1 and CAG3 subunits (Fig 2B and fig. S5D). This reinforces the importance of its activity in photosynthetic organisms (*54*). Complex III is nearly identical to the one from angiosperms, and we observe partial ubiquinone occupancy at the Q_o_ (QH2 oxidation) and Q_i_ (Q reduction) sites (fig. S6). Complex IV, on the other hand, is quite different from what was previously described in angiosperms (*18–20*), yeast (*22*), and animals (*51, 55*). Protein subunits are conserved, but structurally, extensions of Cox2a, Cox2b, Cox5c and Cox6b come together and protrude in the IMS to form what we term an “IMS tail” (fig. S7). Structurally, the IMS tail is similar to what was recently described in Euglena, although the proteins involved are entirely different (*16*).

**Figure 3:**
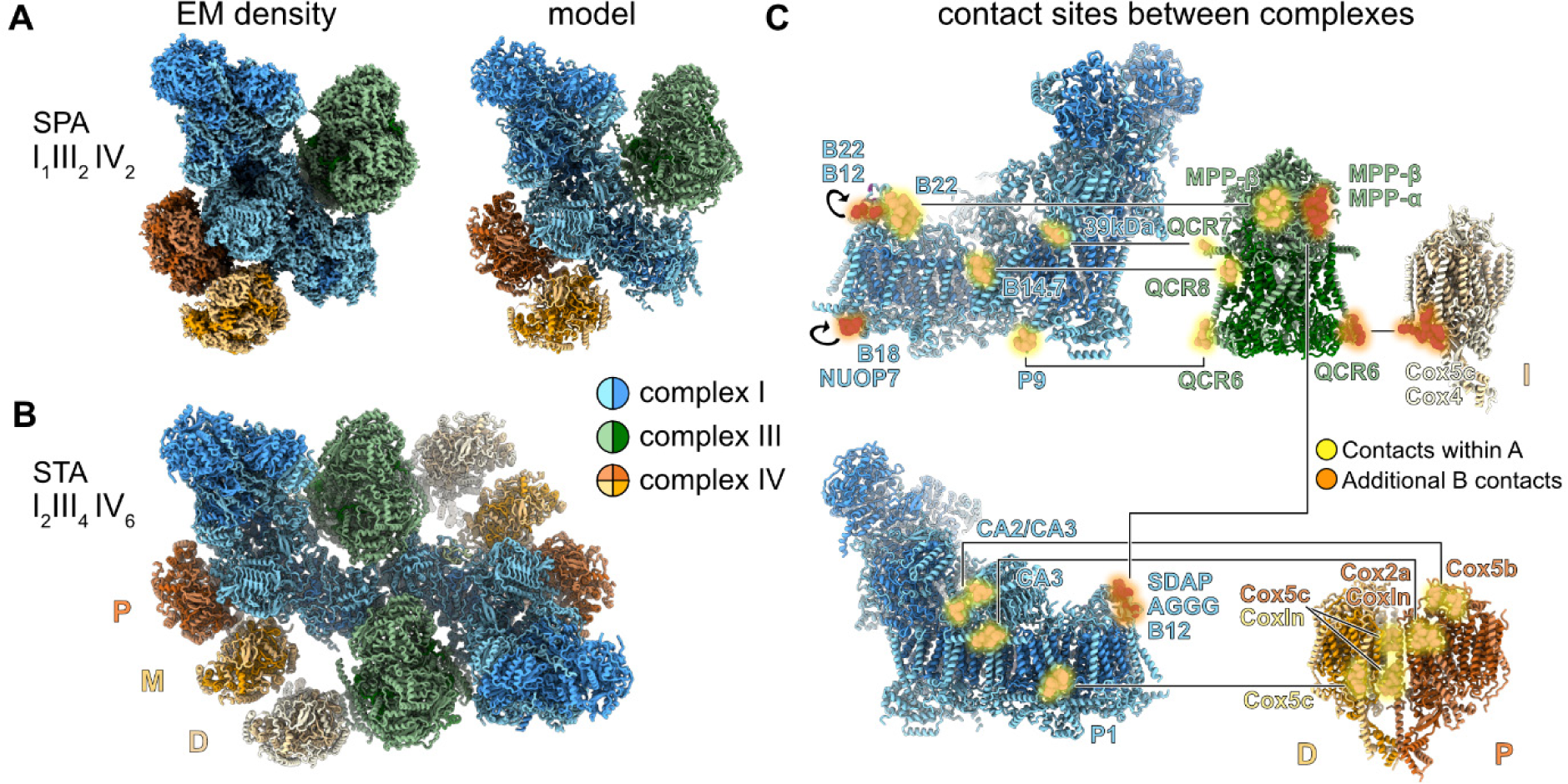
Models of the isolated and native respirasomes. **A)** Single-particle cryo-EM density and resulting molecular model of the isolated I_1_ III_2_ IV_2_ supercomplex. Details of each complex are shown in figs. S5 to S7. **B)** Model of the *in situ* I_2_ III_4_ IV_6_ respirasome, built by combining the STA density (Fig. 2) with the SPA model in panel A. Colored as in Fig. 2. P, M, and D indicate proximal, middle, and distal complex IV monomers. Darker colors: core subunits, lighter colors: supernumerary subunits. **C)** View of the contact sites mediating inter-complex interactions. Yellow: contacts observed both in the isolated and *in situ* structures. Orange: additional contacts observed only in the native I_2_ III_4_ IV_6_ respirasome, mainly mediating dimerization. The names of proteins involved in the contacts are indicated. For B22 and B12, as well as B18 and NUOP7, at the tip of complex I membrane arm, these subunits interact with themselves in the second complex I.

Combining the model obtained from SPA with the STA map of the native respirasome, we were able to build a complete model of the respirasome structure inside the cell (Fig. 3B and Movie 3), which then allowed us to analyze how the different complexes within I_1_ III_2_ IV_2_ (yellow in Fig. 3C) and I_2_ III_4_ IV_6_ (orange in Fig. 3C) interact with each other. Complexes I and III interact via four contacts that are nearly identical to Arabidopsis (*18*). When comparing the ubiquinone binding site, the ubiquinone head in complex I is located about 105 Å from the most proximal Q_o_ and Q_i_ sites in the closest complex III (fig. S8A), which is similar to what was described in the isolated structures of plants (*18*), mammals (*51*), tetrahymena (*17*), and euglena (*16*). This supports the conclusion that the I_1_III_2_ supercomplex is probably the most conserved minimal supercomplex found in mitochondria across eukaryotes. Within I_2_ III_4_ IV_6_, additional contacts are observed that mediate the dimerization of the native supercomplex (shown in orange in Fig. 3C). Notably, the complex III from one asymmetric unit contacts the complex I from the other asymmetric unit. This interaction leads to the complex I monomers being sandwiched on both sides of their distal membrane arm by the two complex III dimers. Additionally, the two complex I monomers contact each other at the distal tips of their membrane arms.

The complex IV monomers in positions P and M interact with each other and complex I, while the monomer in position D interacts with complex III (Fig. 3C). Interestingly, the CoxIn and Cox5c proteins in each complex IV are capable of mediating different types of interfaces either with complex I, IV, or III. This suggests that complex IV may be a more labile component of supercomplexes, contributing to their diversity. This conclusion is in line with recent studies on isolated mouse respirasomes (*12*) and isolated pig mitochondria (*51*), where complex IV binds at different positions around the more structurally conserved I_1_III_2_ core modules.

However, protein-protein interactions alone are likely insufficient to fully stabilize the Chlamydomonas respirasome. We observe only a few inter-complex contacts, compared to the interactions observed in some isolated respirasomes such as the supercomplex from Tetrahymena (*17, 21*). In our structure, we resolve many lipids (figs. S5-S7) stably interacting with proteins that are integral structural components of the complex. From the STA map, we also see that the gaps between the protein complexes of the respirasome are fully filled with lipids. This emphasizes the role of lipids as important structural elements of these supercomplexes (*51*) and might explain why isolating complexes using detergents, which disrupt lipid composition, can result in dissociated and less physiologically relevant complexes. In our case, the distal complex IV was probably lost during biochemical purification because it makes only a few protein-protein interactions that were not sufficient to maintain attachment once the stabilizing lipids were extracted.

### Comparison between isolated and *in situ* structures

Overall, the relative positions of the individual complexes in the SPA map of isolated I_1_ III_2_ IV_4_ and the STA map of native I_2_ III_4_ IV_6_ are very similar. However, there is a notable difference in the activity state of complex I, which suggests that the native complex is active whereas the isolated complex is inactive (see Supplementary text and fig. S9). Moreover, by analyzing the STA map of the native respirasome at a lower threshold, we visualized additional densities attached to the luminal side of the supercomplex, with two densities bound to every complex III dimer and one density bound to every complex IV monomer (Fig. 4A). We could identify these densities as the soluble electron carrier cyt. *c*, with binding sites in agreement with previous *in vitro* data (*56, 57*). The comparably weaker density for cyt. *c* relative the rest of the map likely indicates partial occupancy. This is a good indication that this respirasome is indeed active, as no occupancy or full occupancy might indicate inactive or inhibited states. In total, the I_2_ III_4_ IV_6_ respirasome has four cyt. *c* binding sites on complex III and six binding sites on complex IV, with distances from complex III to complex IV binding sites ranging from ∼90 Å to ∼250 Å (average distance of ∼175 Å). Considering only one respirasome, it would be most favorable for cyt. *c* to travel from complex III to the closest complex IV. However, considering a native crista, we see that respirasomes are located on both membranes, which are separated by an IMS with an average width of ∼150 Å. This means that cyt. *c* could diffuse easily between respirasomes located on both sides of the cristae, and that this would even in some case be more efficient than shuttling within a single respirasome (Fig. 4B).

**Figure 4:**
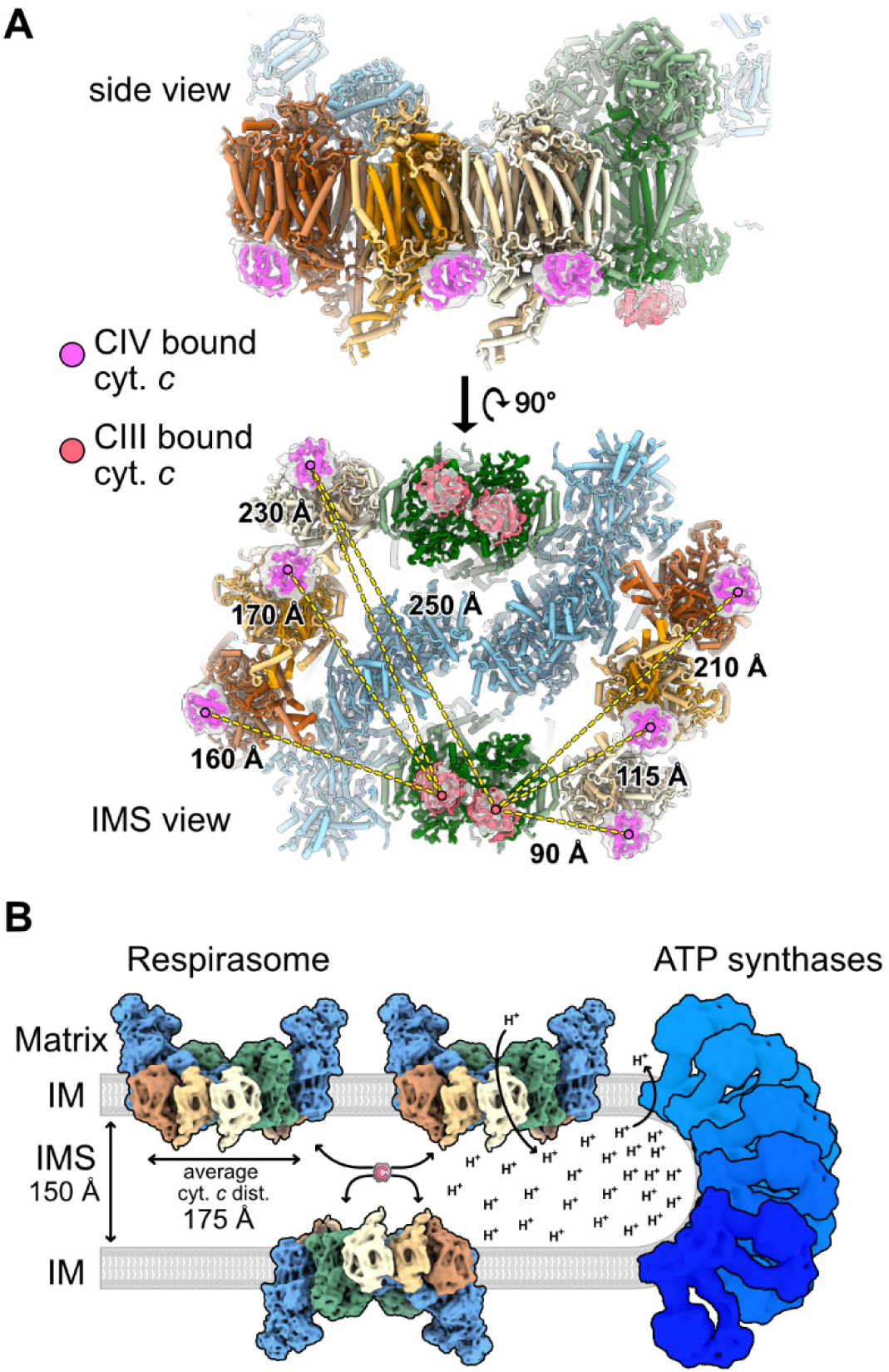
Cytochrome *c* binding sites on the native respirasome. **A)** Views of cyt. *c* densities bound to the respirasome in the STA map, shown from a side view and an IMS view. Salmon: cyt. *c* bound to complex III, magenta: cyt. *c* bound to complex IV. Complex III dimers bind two cyt. *c*, and complex IV monomers bind one cyt. *c*, for a total of ten binding sites in the respirasome. Distances between complex III and complex IV binding sites are indicated, showing the closest (90 Å) and farthest (250 Å) distances for cyt. *c* to travel within the respirasome. **B)** Schematic side view of a crista in *Chlamydomonas reinhardtii*. Proton pumping respiratory complexes are associated together in respirasomes on the flat sides of the crista, while ATP synthases induce high curvature at the crista tip and use the chemical proton gradient to regenerate ATP. The average IMS space (∼150 Å) is shorter than the average distance between cyt. *c* binding sites within a respirasome (∼175 Å), suggesting that cyt. *c* freely diffuses between respirasomes in the crista.

Two models have been put forward to explain the path of electron transfer along the respiratory chain. The substrate channeling model proposes that the electron carriers, ubiquinone and cyt. *c*, are channeled between the different complex I, III, and IV sites of a single respirasome (*58–60*). In contrast, the free diffusion model proposes that the carriers diffuse freely in the IM and IMS, passing between different respirasomes (*58–60*). Considering the overall arrangement and stoichiometry of the Chlamydomonas respirasome, it is possible that reduced ubiquinol from each complex I could be favorably transferred to one of the Q_o_ sites (the proximal site relative to complex I) in a neighboring complex III within the same supercomplex (fig. S8B). However, the position of the other complex III Q_o_ site (distal relative to complex I) appears to be less accessible to ubiquinol. Indeed, it is further away from the complex I Q cavity and faces an enclosed lipid raft, possibly limiting the diffusion of ubiquinol in that area of the supercomplex, even if no sign of symmetry breaking (e.g., locked Rieske subunit (*55*)) could be observed in the structure (fig. S8B). Moreover, densities attributed to cyt. *c* are visible on all complex IV monomers, suggesting they are all active, but only one of these monomers (D) is located adjacent to a complex III. It seems more plausible that electrons are transferred by free diffusion of ubiquinol and cyt. *c* both within a supercomplex and between different respirasomes in the cristae.

### Insights and questions into respirasome function

It remains an open question why respiratory complexes join together into respirasomes (*58*). While roles in respiratory complex activity and assembly have been attributed to supercomplexes (*55, 61–63*), their functional significance remains debated, with substrate channeling in particular being called into question (*58, 63–66*). Recent studies in mice and Drosophila, which biochemical assays show harbor decreased levels of supercomplexes, indicate that respirasomes may not be required for normal respiration and physiology (*67, 68*). Nevertheless, evolution appears to have repeatedly selected for respirasomes, as many of the additional proteins acquired by respiratory complexes in different eukaryotic lineages serve to mediate the formation of these supercomplexes (*20, 58, 60*).

Our study of native respirasomes inside Chlamydomonas cells provides new clues into the functional significance of respirasomes. As described above, we find only minor support for the hypothesis of substrate channeling (perhaps of ubiquinol from complex I to III) and conclude that respirasomes likely do not accelerate the free diffusion of cyt. *c* from complex III to IV. A second hypothesis for respirasome function is that they may serve to control respiratory complex stoichiometry, but this is challenged by the many different supercomplex stoichiometries observed to date, sometimes even within the same species (*16, 20, 51, 58, 60*). Here in Chlamydomonas, we describe the new stoichiometry of a I_2_ III_4_ IV_6_ respirasome. It is not obvious why this particular stoichiometry is favored, but perhaps the relative excess of complex IV might help ensure efficient capture of cyt. *c* and minimize the ROS production by consuming the O_2_ in the vicinity of complex III. A third hypothesis for respirasome function is that the supercomplex interactions may help stabilize assembly intermediates of respiratory complexes, aiding in their biogenesis (*55, 61–63*). Our data is consistent with this idea, as we do not see free complex I or complex III, and we observe a minor fraction of respirasomes at the IM, where complex biogenesis occurs. A final hypothesis is that respirasomes function to shape crista membranes, with ATP synthase oligomerization directing high-curvature folding of the crista tips and respirasomes shaping the surface of the cristae (which is flat, with the exception of tubular Tetrahymena cristae shaped by curved respirasomes (*17*)). As visualized by *in situ* cryo-ET (Fig. 1D), this membrane architecture creates a narrow luminal space and lateral heterogeneity with the cristae, thereby generating a gradient of proton flux from respirasome source to ATP synthase sink (Fig. 4B) (*69*). In such a manner, respirasomes would enable efficient respiration through the indirect mechanism of establishing crista architecture and molecular organization. The native respirasome structure presented in our study provides a blueprint to specifically disrupt supercomplex formation *in vivo* and mechanistically dissect the physiological relevance of these enigmatic molecular machines.

## Supplementary Figures

**Supplementary Fig. 1:**
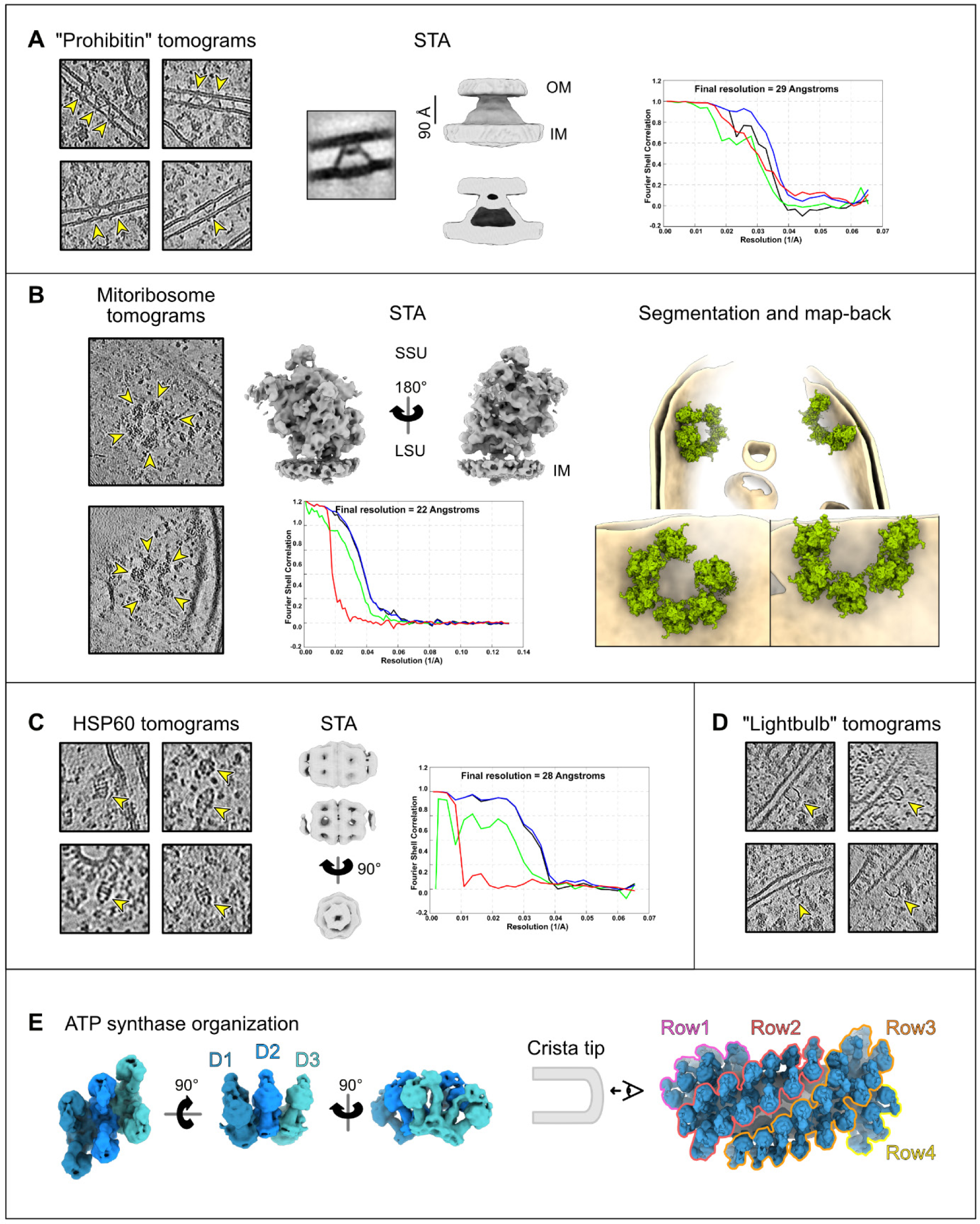
Snapshots, STA, and organization of complexes resolved in native mitochondria. A) Zoomed-in views of denoised tomograms containing putative prohibitins that were used for STA. The resulting STA map and FSC plot are shown alongside a view of a slice through the volume from IMOD. B) Zoomed-in views of denoised tomograms containing membrane-bound mitoribosomes organized as circular polysomes. The STA map and FSC plot are shown with segmentation and map-back of a mitochondrion containing circular mitoribosomal polysomes. C) Zoomed-in views of denoised tomograms containing HSP60; side and top views are shown. The STA map at high and low threshold and FSC plot are shown. D) Zoomed-in views of denoised tomograms containing the lightbulb-shaped unknown complex, located on the mitochondrial OM. E) View of ATP synthases organization. On the left, the basic organization of dimers in row are shown, with each ATP synthase dimer colored in a different shade of blue. On the right, the higher-order organization of the ATP synthase rows is shown. Several rows are stacked on the cristae tip to induce the full fold of the membrane.

**Supplementary Fig. 2:**
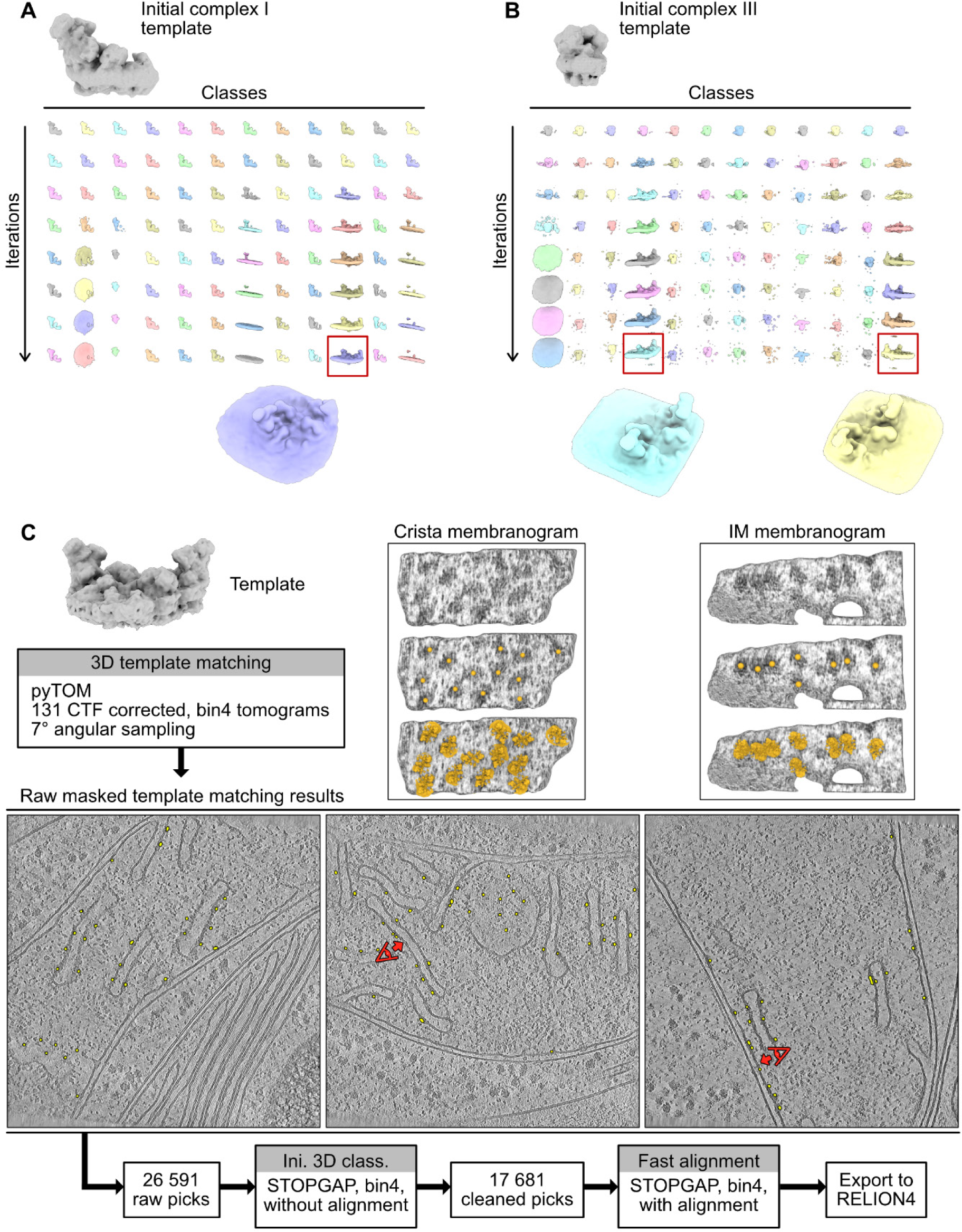
STA processing: template matching and initial identification of respirasomes. Graphical summary of the processing workflow described in Methods. A and B) initial identification of the respirasomes using either a complex I reference (A) or a complex III reference (B). After classification, all good particles for both complex I and complex III converged to respirasomes (magnified view of the classes shown). C) Template matching using the respirasome template derived from A and B. Raw template matching cross-correlation peaks (yellow) are overlayed on 3 tomograms for illustration. Membranograms (*70*) of a crista and an inner membrane are shown. Center picked positions obtained by template matching after initial cleaning are shown in orange (middle), and a view with the template overlayed is also shown (bottom). A red eye with an arrow indicates which membranes were used for the membranograms above.

**Supplementary Fig. 3:**
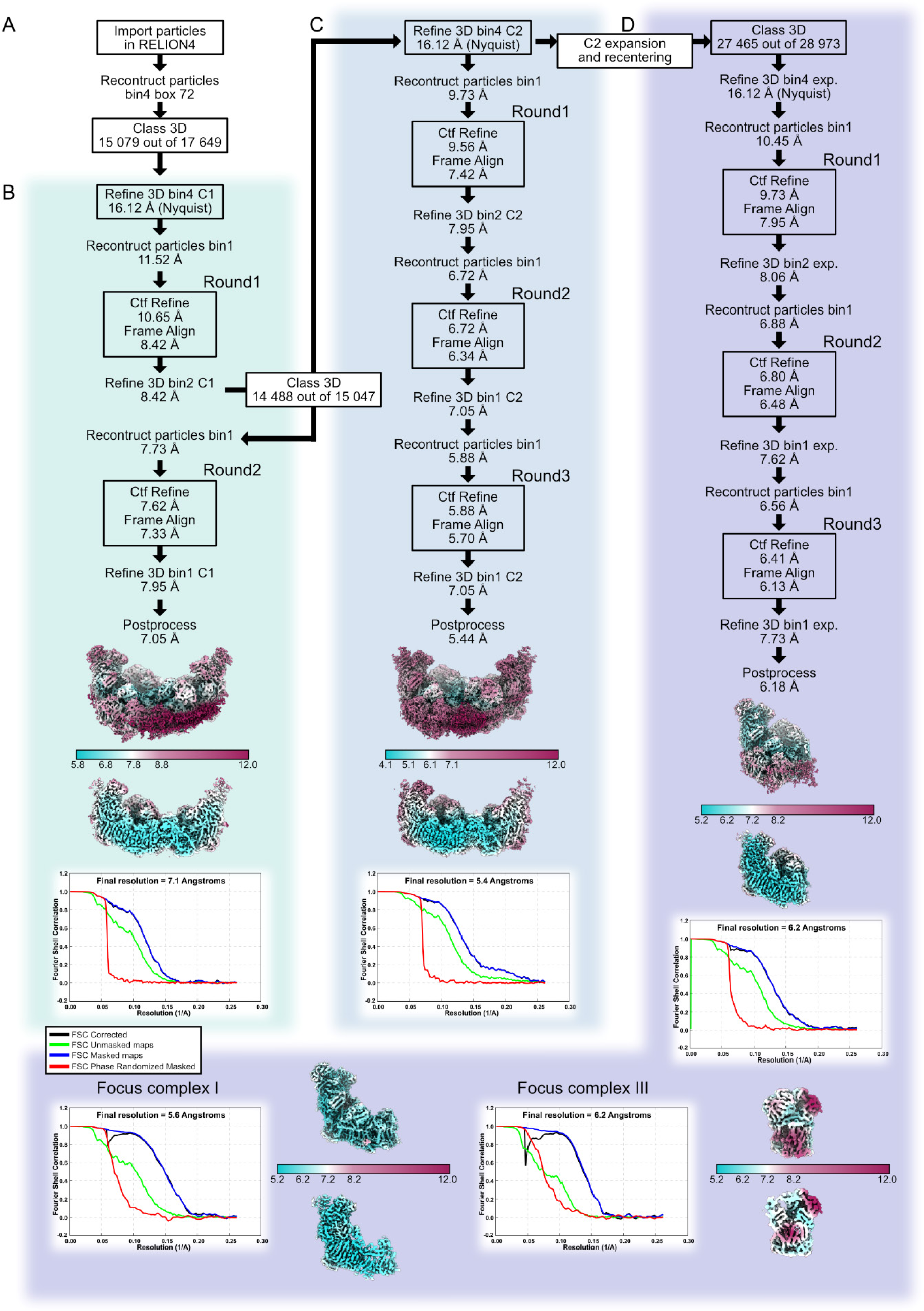
STA processing: high-resolution refinement. Graphical summary of the processing workflow described in Methods. A) import to RELION-4 and further cleaning of the particles. B) Refinement without symmetry (C1 symmetry). C) Refinement with C2 symmetry. D) Refinement for C2 symmetry-expanded particles. For the final reconstruction, “gold standard” FSC (GSFSC) curves are plotted, with resolution calculated at the 0.143 threshold. The estimated local resolution plotted on the maps were generated in RELION-4, with maps also shown in cut view.

**Supplementary Fig. 4:**
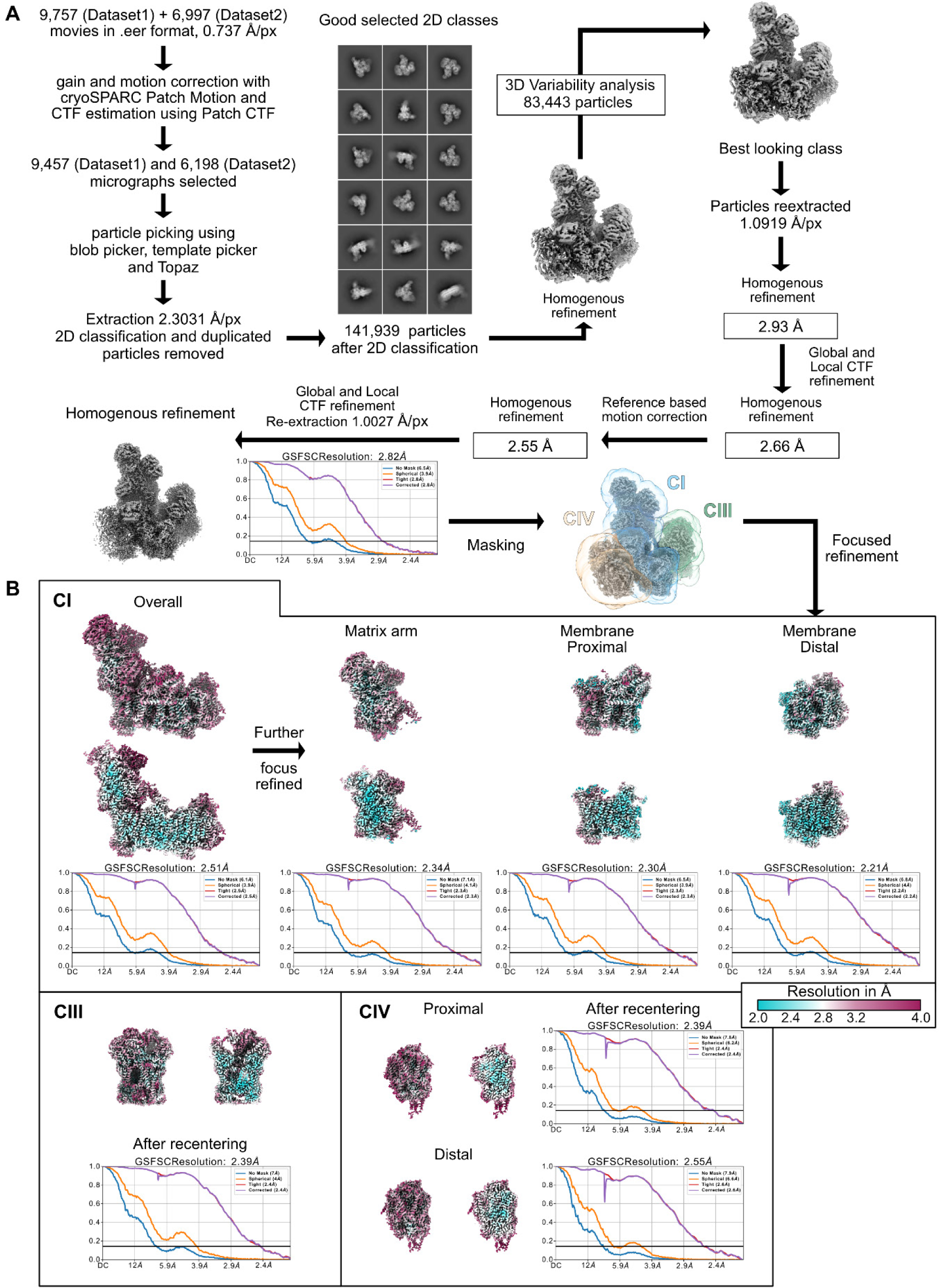
Single-particle data processing workflow. Graphical summary of the processing workflow described in Methods. A) Post-processing, 2D and 3D classification. B) Focused refinements. For the final reconstruction, GSFSC curves are plotted, with resolution calculated at the 0.143 threshold. Local resolution plotted on the maps were generated using the built-in cryoSPARC tool with default parameters (FSC=0.5 threshold), all using the same resolution scale, with maps also shown in cut view.

**Supplementary Fig. 5:**
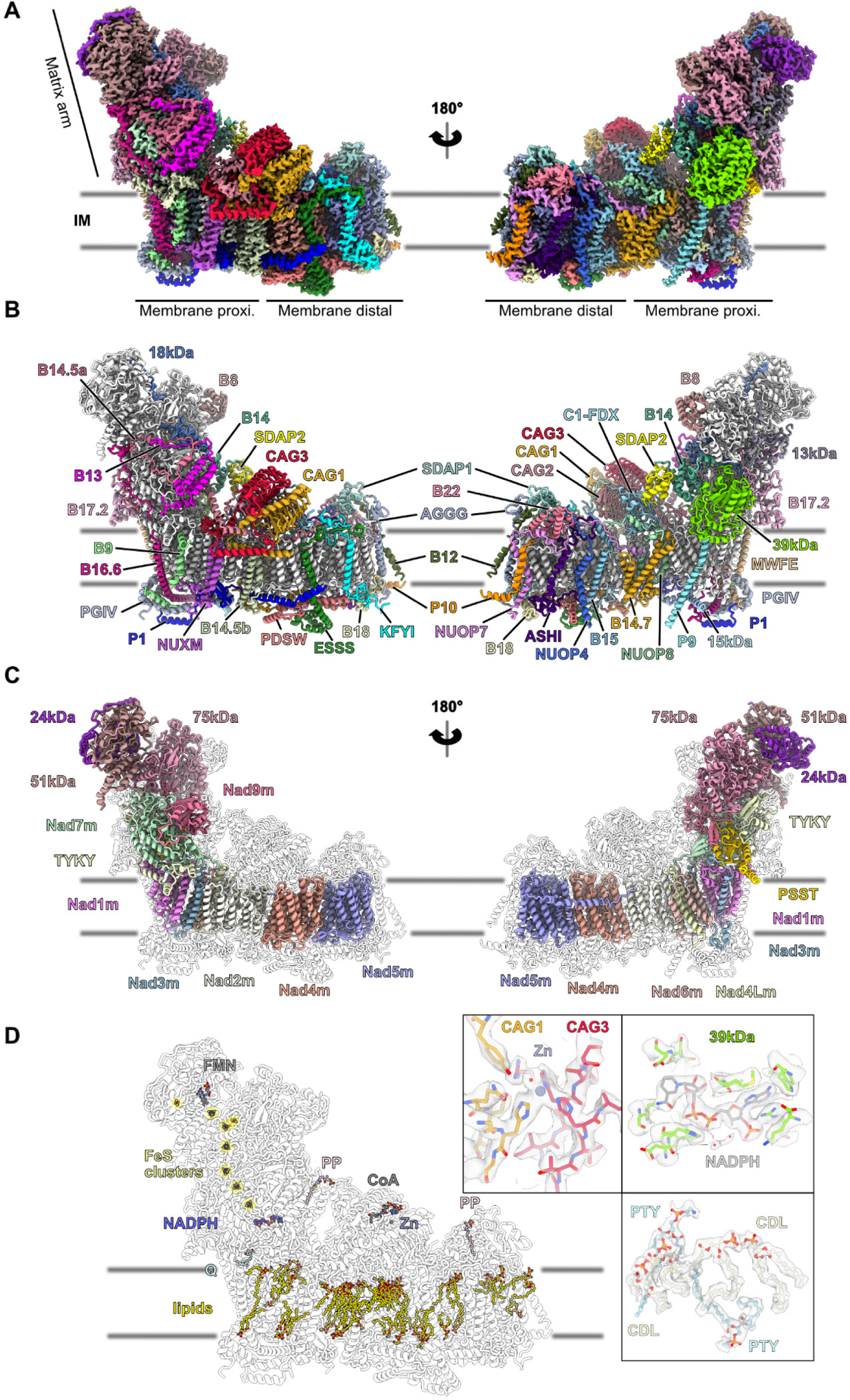
Details of *C. reinhardtii* complex I. **A)** Composite single-particle cryo-EM map of Chlamydomonas complex I. Each of the 51 proteins is depicted with a different color. **B and C)** The resulting model, depicting the 37 annotated supernumerary subunits (B) and the 14 core subunits (C). IM: inner membrane. **D)** View of the different resolved ligands, with lipids in the membrane arm shown in yellow and iron sulfur clusters in the matrix arm highlighted with a yellow halo. FMN: flavin mononucleotide in the 51kDa subunit, PP: S-acyl-4’-phosphopantetheine in the SDAP subunits. Contrary to Arabidopsis and like Polytomella, the bridge subunit C1-FDX is inactive and does not bind an iron atom. Similarly, no zinc is bound to the 13kDa subunit. Partial density was observed for the ubiquinone (head group not fully resolved), as well as partial density for CoA in the carbonic anhydrase. Close-up views with density overlayed are shown for the active site in the carbonic anhydrase domain, NADPH in the 39kDa subunit, and of phosphatidylethanolamine (PTY) and cardiolipins (CDL) next to the B14.7 subunit.

**Supplementary Fig. 6:**
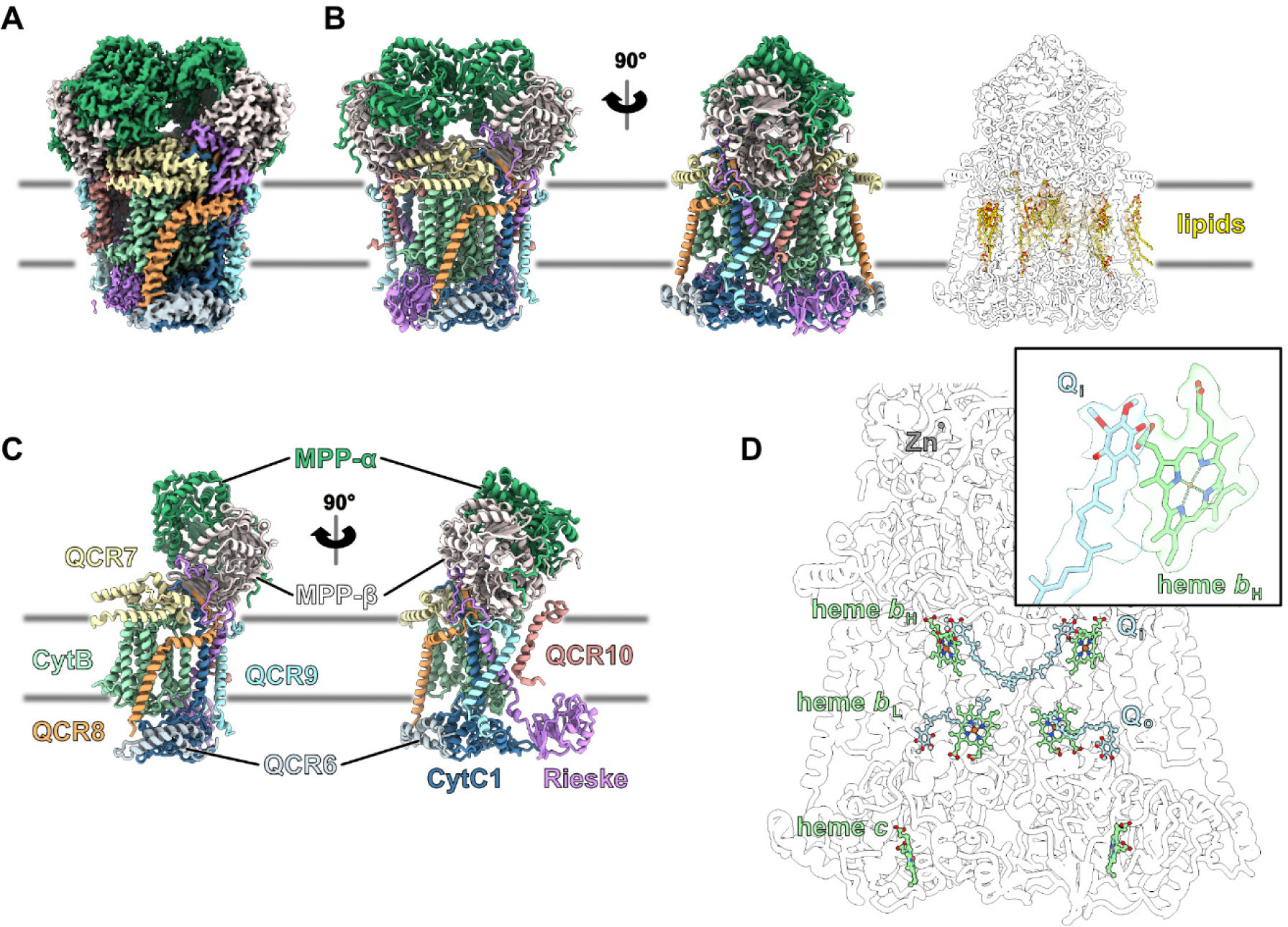
Details of *C. reinhardtii* complex III. **A)** Single-particle cryo-EM map of Chlamydomonas complex III. Each of the 10 proteins is depicted with a different color. The Rieske protein was only partially resolved due to high flexibility. **B)** The resulting model, with lipids (yellow) shown in the rightmost picture. **C)** The annotated 10 proteins within a monomer. **D)** Heme groups (green) and ubiquinol sites (blue) are shown, along a magnified view of the fully resolved ubiquinol next to one of the hemes *b*_H_ in their density (partial density of the head group is observed for the 3 other sites). Abbreviations: Q_i_ Q reduction; Q_o_ Q oxidation; b_H_ high-potential heme b; b_L_ low potential heme b.

**Supplementary Fig. 7:**
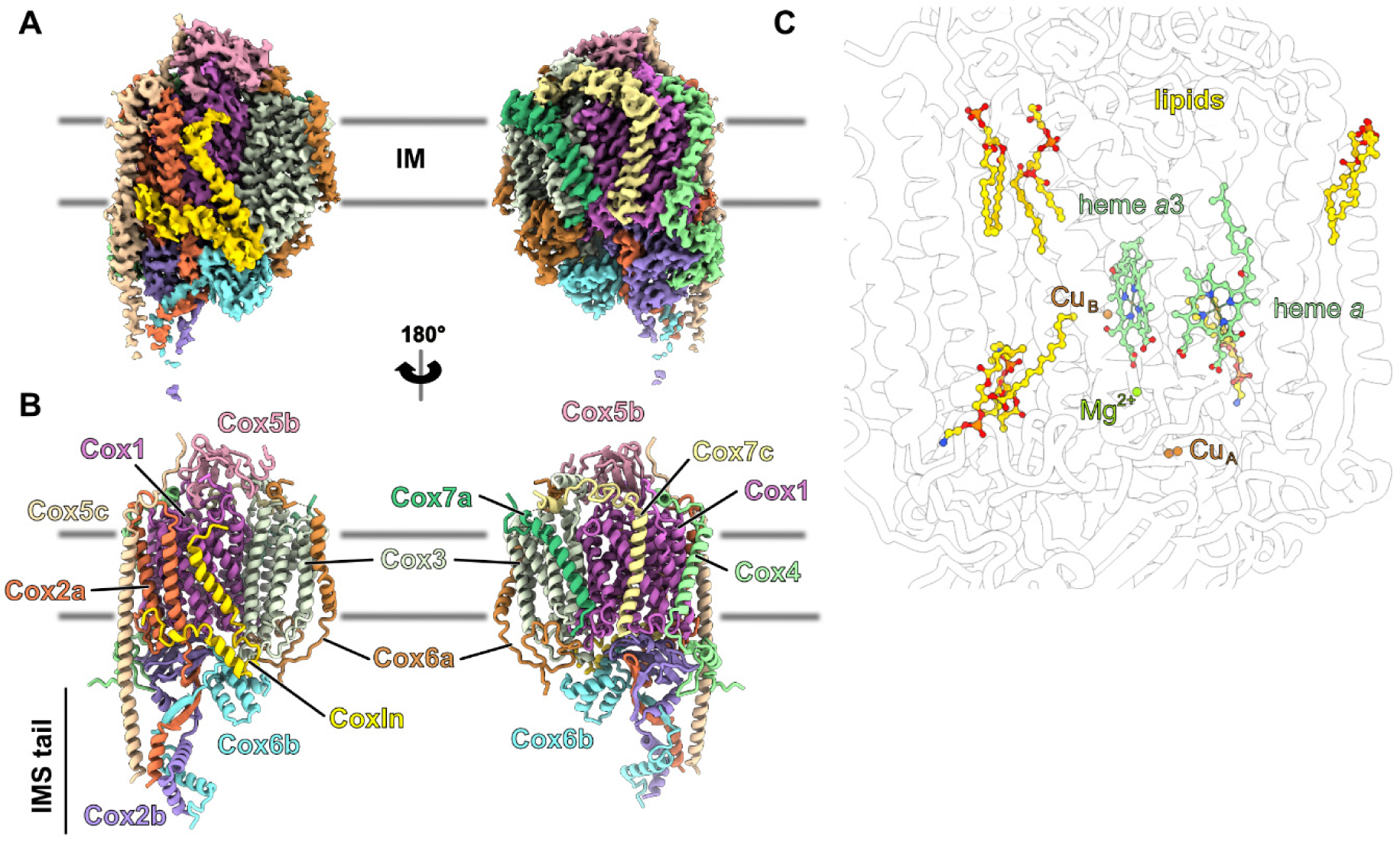
Details of *C. reinhardtii* complex IV. A) Single-particle cryo-EM map of Chlamydomonas complex IV. Each of the 14 proteins is depicted with a different color. B) The resulting model with the annotated protein subunits. Extensions of Cox2a, Cox2b, Cox5c and Cox6b come together to form the IMS tail and protrudes in the IMS. C) View of the lipids (yellow), the heme groups, and the copper co-factors.

**Supplementary Fig. 8:**
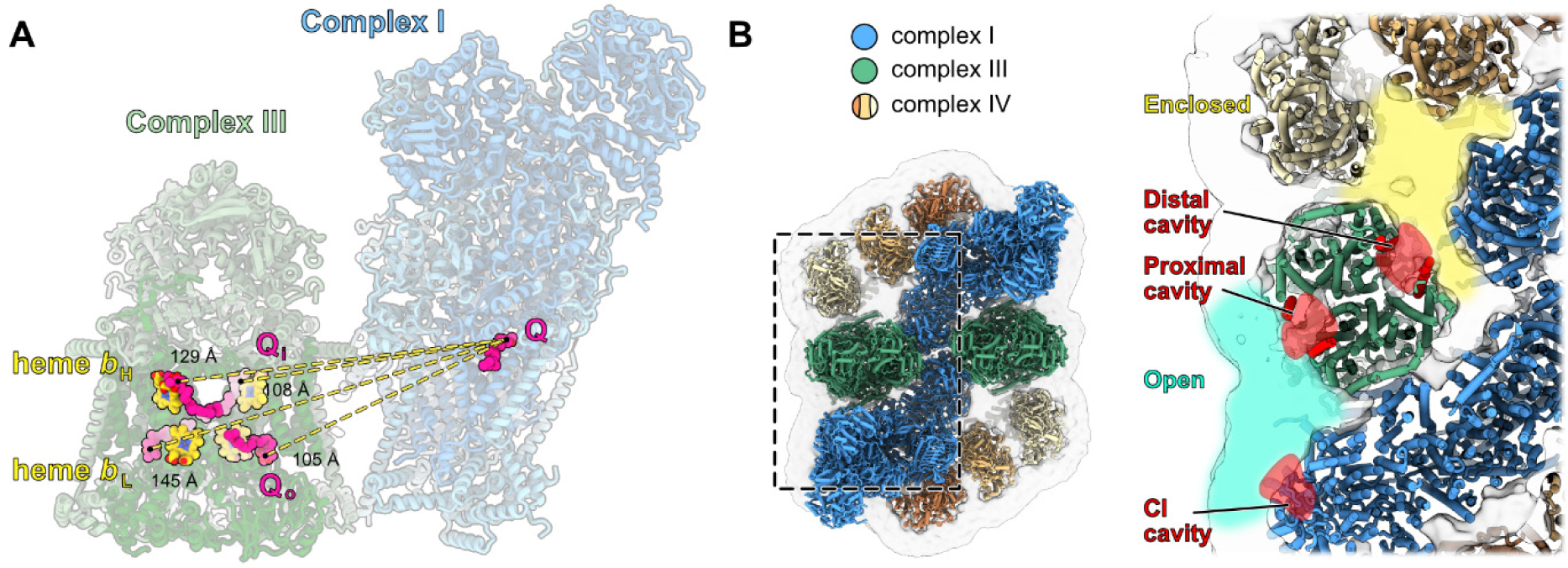
Complex I and complex III Q sites. A) Distances from the complex I Q site to the two Qi and Qo sites in complex III. B) Top view of the native respirasome and zoomed-in cut view of complexes III and I. The complex I Q cavity is open to the membrane (blue) and closest to one of the CIII cavities (proximal). The other CIII cavity (distal) is facing an almost closed membrane lipid raft delimited by CI, CIII, and two CIV.

**Supplementary Fig. 9:**
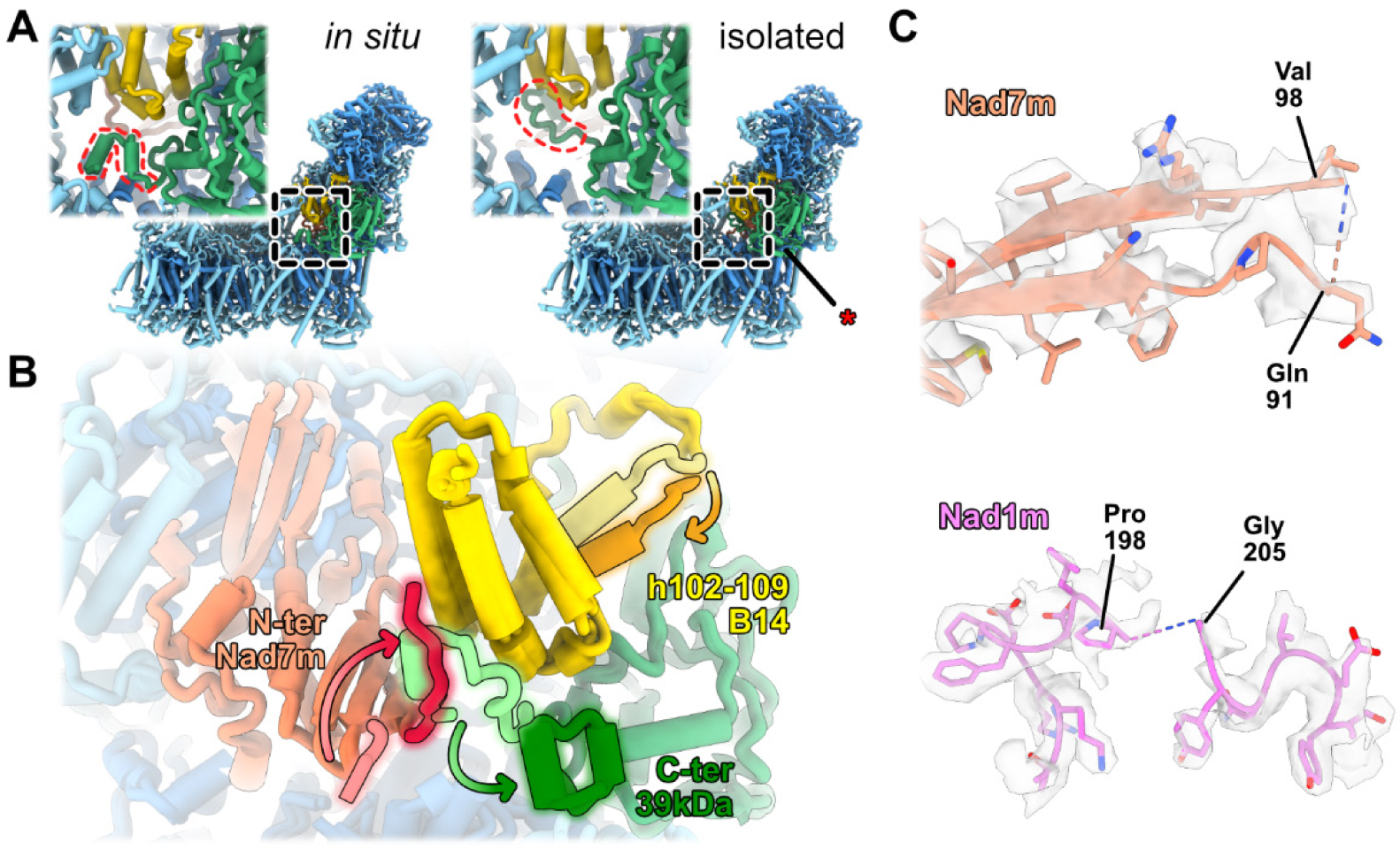
Conformational differences in complex I between native and isolated structures. A) Different conformations of the C-ter of the 39kDa subunit observed for complex I between the isolated and *in situ* (in-cell) structures. The red asterisk marks the location of the disordered Nad1m and Nad7m. B) Magnified view of the rearrangement of the B14, Nad7m, and 39kDa subunits of complex I between the *in situ* and isolated structures. Positions of the rearranged proteins are showed, with lighter colors corresponding to the isolated conformation and darker colors to the *in situ* conformation. In the isolated conformation, the N-ter of Nad7m (pink) extends away from B14 and does not contact other proteins, while the C-ter of 39kDa (light green) is bound to B14. In the *in situ* conformation, the C-ter of 39kDa (dark green) releases B14, while the N-ter of Nad7m (red) contacts B14. Additionally, the helix formed by residues 102-109 (termed h102-109) of B14 switches from an upward position in the isolated structure to a downward position in the *in situ* conformation. C) View of the disordered loops of Nad1m and Nad7m.

## Supplementary Tables

**Supplementary Table 1:**
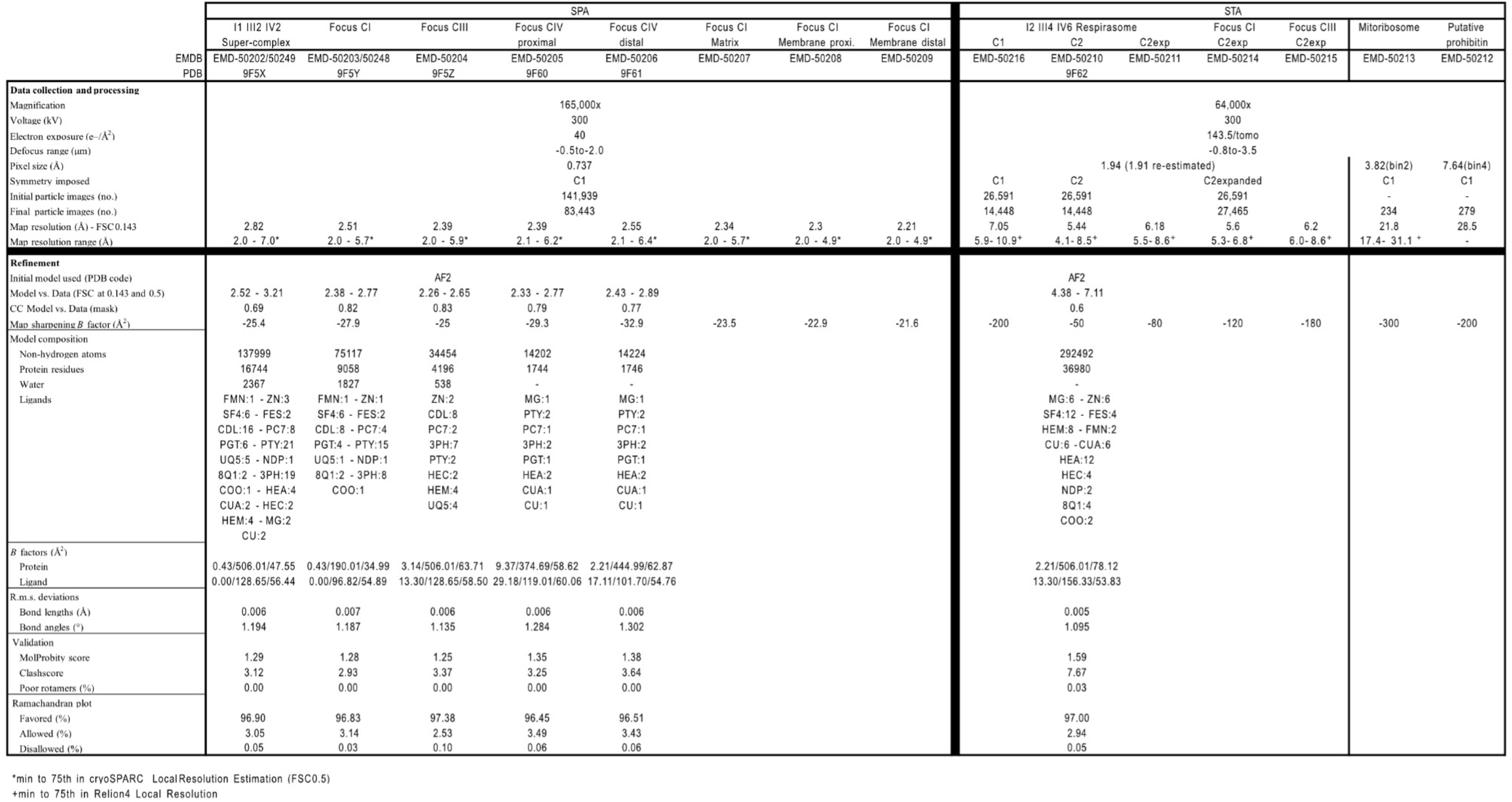
Cryo-EM validation.

**Supplementary Table 2:**
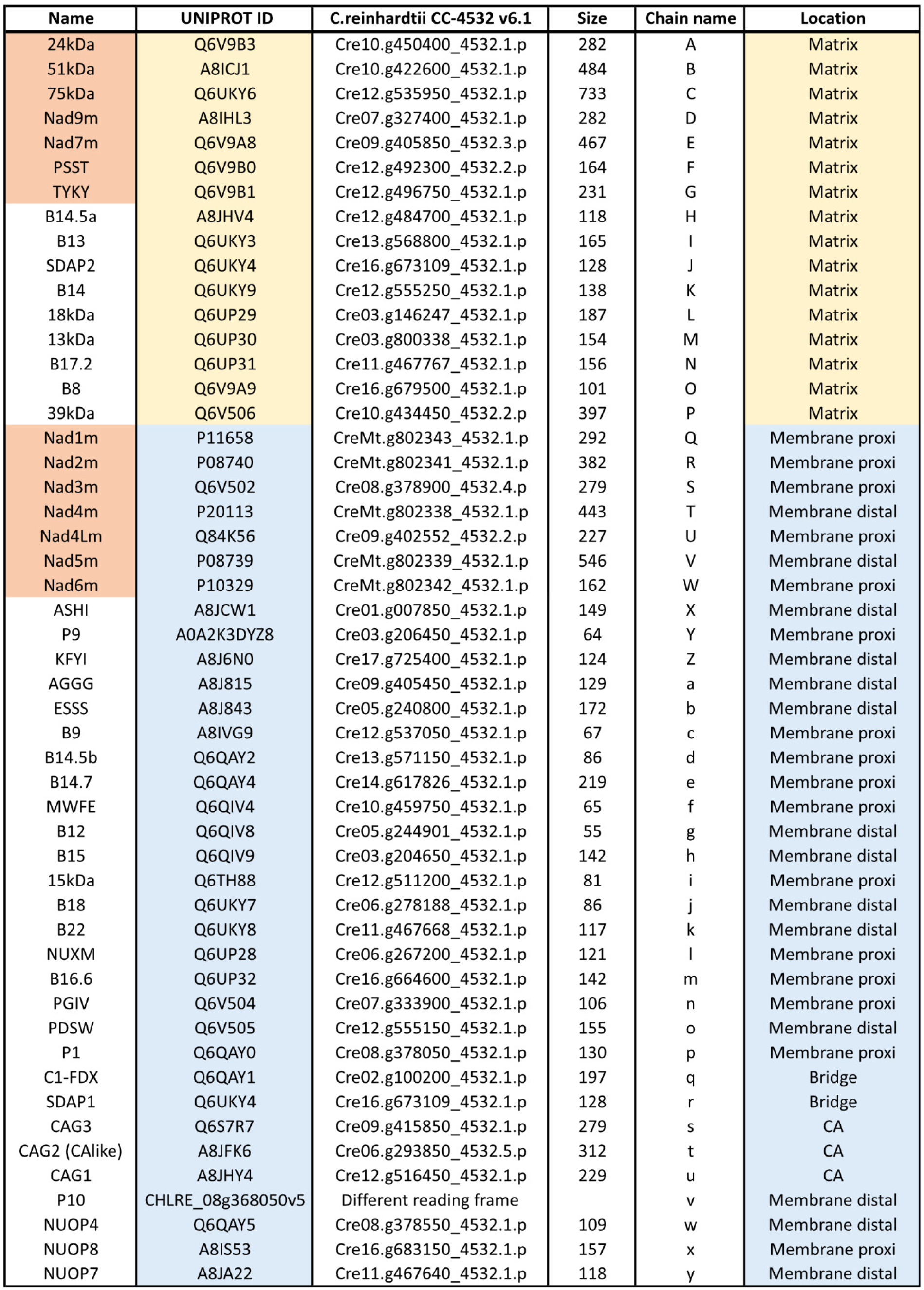
List of complex I proteins. List of the protein subunits constituting *C. reinhardtii* respiratory complex I. The table is divided between matrix arm proteins (yellow) and membrane arm proteins (blue). The names of the 14 core subunits are colored in red.

**Supplementary Table 3:**
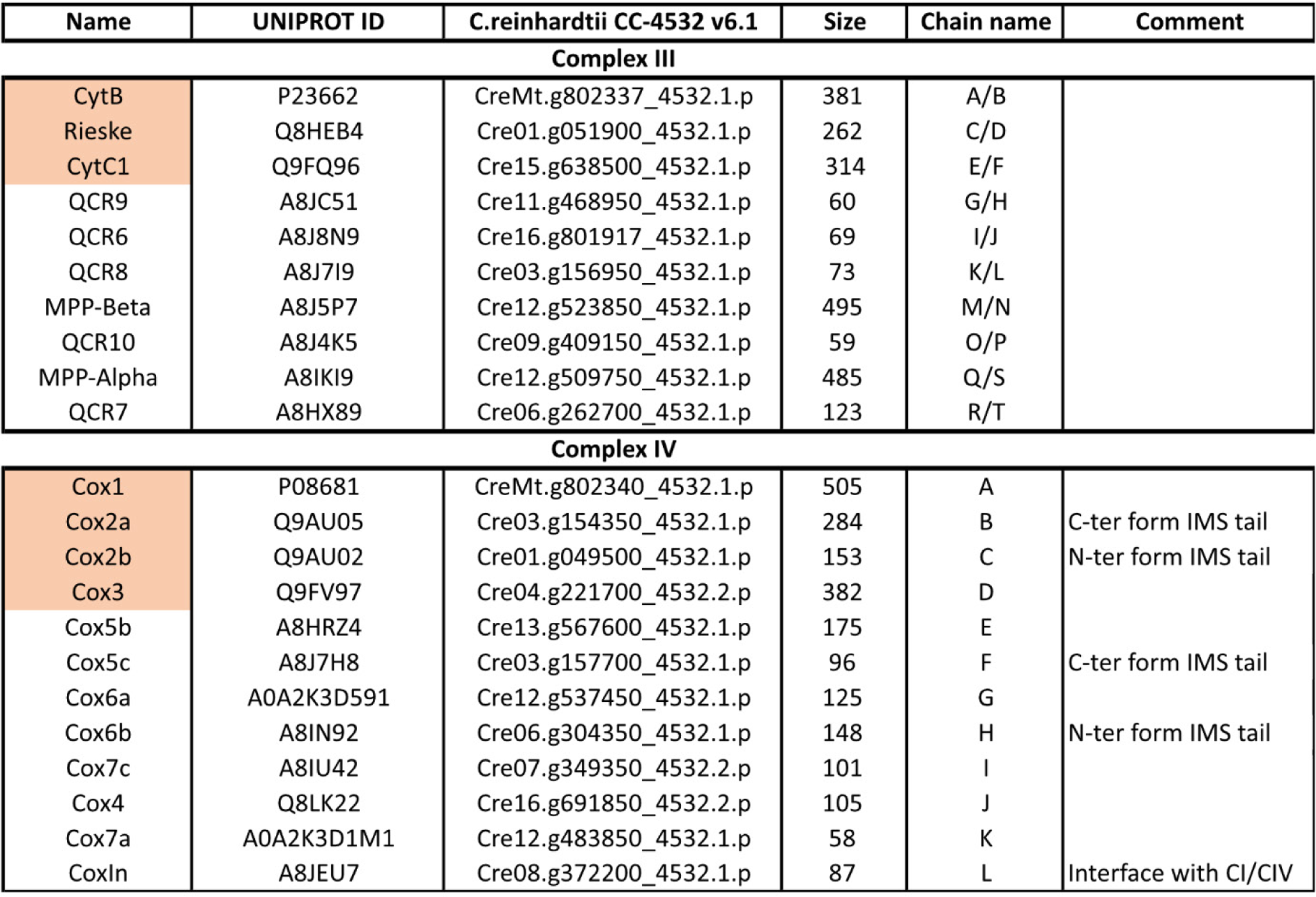
List of complex III and IV proteins. List of the protein subunits constituting *C. reinhardtii* respiratory complexes III and IV. The names of the core subunits (3 in complex III, 4 in complex IV) are colored in red.

## Supplementary Movie Legends

**<u>Movie 1:</u>** Tomogram and map-backs

Sequential sections back and forth through a tomographic volume, followed by a reveal and tour of a mitochondrion, showcasing its architecture and the different molecular complexes identified.

**<u>Movie 2:</u>** In cell subtomogram average of the respirasome

The movie tilts from top side to top view, and shows the different components of the Chlamydomonas respirasome in native cells. The respirasome rotates to show the bended membrane. Finally, individual complexes I, III and IV are shown along with zoom-in detailed views of the densities.

**<u>Movie 3:</u>** Single-particle structure and resulting models

The movie shows the single-particle cryo-EM map of the I_1_III_2_IV_2_ purified supercomplex. It rotates, and shows the resulting atomic model. Then the atomic model of I_2_III_4_IV_6,_ built from the native density and the model obtained from single-particle, is shown.

**<u>Movie 4:</u>** Morph between the isolated and in cell conformations of complex I

The movie shows the rearrangements of the complex I subunits Nad7m, B14 and 39kDa between the isolated and native complexes

## Supplementary Text

### Protein-protein interactions within the supercomplexes

Complexes I and III interact via four contacts: the tip of P9 with QCR9, B14.7 with QCR8, 39kDa with QCR7 and B22 with MPP-β (shown in yellow in Fig. 3C). These contacts are nearly identical to those observed in Arabidopsis (*18*), apart from the additional 39kDa - QCR7 contact point.

On the other side of complex I, two complex IV monomers are found. The interactions are mediated by the CA2 and CA3 subunits of the carbonic anhydrase module in the complex I with Cox5b, Cox2a and CoxIn of the proximal complex IV. In the distal complex IV, its subunit Cox5c interacts with the P1 protein of complex I. Finally, the proximal and middle complex IV interact with each other via the Cox5c and CoxIn subunits. Although no direct contacts are observed between the IMS tails of the proximal and middle complex IV, considering the short distance separating them, it cannot be excluded that they might interact via the IMS tails. Also to be noted, in the I_1_ III_2_ IV_2_ supercomplex, no direct contact is formed between complexes III and IV.

Within I_2_ III_4_ IV_6_, additional contacts are observed that mediate the dimerization of the supercomplex (shown in orange in Fig. 3C). Notably, complex III from one asymmetric unit forms contacts with the complex I of the other asymmetric unit. This interaction is mediated by subunits SDAP, AGGG and B12 in complex I with MPP-β and MPP-α in complex III. The distal membrane arms of complex I from each asymmetric unit contact each other the via subunits B22 and B12 on the matrix side, as well as B18 and NUOP7 on the IMS side. Finally, the distal complex IV, which is only present in the native structure, forms direct contacts via Cox5c and Cox4 with protein QCR6 of complex III from the other asymmetric unit. This is the only interaction that this complex IV makes with other complexes within the supercomplex, likely explaining its high structural heterogeneity.

### Conformational differences in complex I between native and isolated structures

Complex I adopts a different conformation between the purified and native complexes (fig. S9 and Movie 4). At the junction between the matrix arm and proximal membrane region, at the base of the bridge domain, the C-ter part of the 39kDa subunit is largely shifted between the SPA and the STA structures. Further inspection allowed us to observe rearrangements also in subunits B14 and Nad7m (fig. S9). Indeed, one helix (h102 – 109) of B14 adopts a shifted position, and the C-ter part of the 39kDa subunit is also positioned differently between the SPA and the STA structures. In the SPA structure, the C-ter part of the 39kDa protein extends and binds B14. However, in the STA map, C-ter part of the 39kDa subunit is completely shifted toward Nad6m, and the position initially occupied by 39kDa is instead occupied by the N-ter of Nad7m.

Complex I activity is performed through the so-called coupling mechanism, which involves coordination between electron transfer from NADH to ubiquinone in the matrix arm, and proton translocation in the membrane arm (*71, 72*). This is facilitated by conformational changes, notably hinging of the matrix arm relative to the membrane arm (*73, 74*). The changes observed here might be a way to regulate complex I activity, with C-ter part of the 39kDa subunit preventing the hinging of the matrix arm relative to the membrane arm, locking the complex in a specific conformation. Further inspection of Nad1m and Nad7m in the SPA maps revealed faint densities for two flexible loops, which are indicators of complex I state. Here, the Nad1m loop (198 – 205aa) and Nad7m loop (91 – 98aa) are disordered (fig. S9), which is a hallmark of a deactivated complex I in mammals (*51, 71*). Thus, it appears that the SPA structure represents a mix of several states where complex I is mostly deactivated, whereas in the native STA map, complex I is mostly in an active conformation. Considering that the rearrangements of the 39kDa, B14, and Nad7m subunits between the in-cell and isolated complexes involve helices and extensions only found in chlorophytes, this hints at a specific way of regulating the complex I coupling mechanism.

## Material and Methods

### Cryo-ET data acquisition and preprocessing

All data analysis was performed on the cryo-ET dataset deposited under EMPIAR-11830. Briefly, tilt-series data were collected using a Titan Krios G4 transmission electron microscope operating at 300 kV equipped with a Selectris X energy filter with a slit set to 10 eV and a Falcon 4i direct electron detector (Thermo Fisher Scientific) recording dose-fractionated movies in EER format. A dose-symmetric tilt scheme using TEM Tomography 5 software (Thermo Fisher Scientific) was employed in the acquisition with a tilt span of ± 60°, covered by 2° or 3° steps starting at either ± 10° to compensate for the lamella milling angle. Target focus was set for each tilt-series in a range of -1.5 µm to -3.5 µm in steps of 0.25 µm. The microscope was set to a magnification corresponding to a pixel size of 1.96 Å at the sample level and a nominal dose of 3.5 e^-^/Å^2^ per tilt image.

The tilt series data were then preprocessed using TOMOMAN (*75*) for streamlining all the following operations. Raw EER data were motion-corrected using RELION’s implementation of MotionCor2 (*76*). Defocus estimation was carried out using the TiltCTF (*75*) procedure in TOMOMAN based on CTFFIND4 (*77*). Dose weighting was performed internally by TOMOMAN. Fiducial-less tilt series alignment was carried out using AreTomo (*78*), and tomograms were reconstructed using IMOD (*79*) and novaCTF (*80*). Denoising was performed with cryo-CARE on tomogram pairs reconstructed from odd or even raw frames respectively (*81*).

### Subtomogram averaging

Processing workflow is shown in figs. S2 and S3. To start, 131 mitochondria tomograms from the Chlamydomonas dataset, EMPIAR-11830, were selected.

Initial detection of respirasomes were done using complex I and complex III models lowpass filtered to 16 Å and used for template matching in STOPGAP (*82*) on a subset of 15 CTF corrected tomograms. After extraction and classification of the picks in STOGAP, the totality (at this step) of the complex I and complex III good particles converged to I_2_ III_4_ IV_6_ respirasomes (fig. S2A-B). The recentered low-resolution average of the respirasome was then used for template matching. To pick the particles, 3D template matching was performed on bin4 tomograms (7.84 Å /pixel) with a 7° angular sampling using pyTOM (*83, 84*) and STOPGAP (*82*) (fig. S2C). Picks were extracted using a mask covering only mitochondria volumes to exclude template-matching hits outside of mitochondria, hence limiting obvious false positives. For each tomogram, the number of picked particles were thresholded so that each tomogram would be largely over-picked, to maximize the number of true positives, yielding an initial set of 26,591 candidate particle picks. These picks were then imported into STOPGAP (*82*) and the particles were cleaned by several rounds of 3D classification without alignment using bin4 subtomograms of box size 64, to clean out obvious false positives (membranes and high-signal contaminants). At this step, 17,681 particles were left. The remaining particles were then exported to RELION-4 (*85*) for high resolution refinement (see fig. S3). Bin4 subtomograms were generated with a box of 72 pixels and further cleaned in 3D classification, 15,079 particles were left. Particles were first 3D refined in C1 at bin4 and reached Nyquist, 16.12 Å. A first cycle of CtfRefine and FrameAlign allowed reaching 8.42 Å. From there particles were further 3D classified and 14,488 particles were kept. These particles were used for further processing in C1 or with C2 symmetry. In C1, a second round of CtfRefine and FrameAlign jobs, followed by 3D refine at bin1, reached 7.05 Å. In parallel, particles were 3D refined in C2 at bin4 and reached Nyquist as well. These particles were used for further processing in C2 symmetry or using symmetry expansion (*86*). Particles treated with C2 symmetry were submitted to three iterative rounds of CtfRefine, FrameAlign and 3D refine which led to resolution going from 9.73 Å before, to 6.72 Å, 5.88 Å and finally 5.44 Å after each round. Particles C2 expanded were recentered on one of the units and 3D classified, resulting in 27,465 particles remaining. Like C2 processing, 3 iterative rounds of CtfRefine, FrameAlign and 3D refine were performed, which led to resolution going from 10.45 Å before, to 6.88 Å, 6.56 Å and finally 6.18 Å after each round. Finally, focused refinement on complex I reached 5.6 Å, and complex III reached 6.2 Å. Note that the entire processing was performed with a pixel size of 1.94 Å /pixel, but post-processing was performed with a re-estimated pixel size of 1.91 Å /pixel.

For mitochondrial ribosomes, a template was generated using the PDB models 7PKQ and 7PKT and low pass filtered to 20 Å and then used for 3D template matching using pyTOM exactly as described for the respirasomes. Only 39 tomograms contained mitoribosomes. Given the low abundance of mitoribosomes, picks were manually curated resulting in 234 final particles which were first quickly aligned in STOGAP, and then imported in RELION-4 for 3D refinement, CTF refinement and FrameAlign, where a similar procedure to respirasome was applied. The final average reached 21.8 Å.

For the putative prohibitin complexes, template matching approaches failed. Hence, 279 particles were manually picked from 33 tomograms. Particles were then imported and aligned in STOPGAP. Final average reached 28.5 Å in resolution. A similar approach was used for HSP60. 138 particles were manually picked from 16 tomograms. Particles were then imported and aligned in STOPGAP, initially without symmetry, and then with D7 symmetry. Final average reached 28 Å in resolution.

### Single particle cryo-EM sample preparation

Mitochondria were purified from *Chlamydomonas reinhardtii* cell wall-less strain CC-4351 as previously (*38*). Cells were grown in Tris-Acetate Phosphate (TAP) medium, under continuous white light (50 µE.m^-2^.s^-1^). Cells were harvested by centrifugation 10 minutes 1000 g, and resuspended in ice-cold Lysis Buffer (25 mM phosphate buffer pH 6.5, 6 % PEG 6,000, 0.5 % (w/v) bovine serum albumin (BSA), and 0.016 % (w/v) digitonin) to a final concentration of 3x10^8^ cells/ml. The suspension was warmed rapidly to 30°C, shaken for 30 seconds, and immediately cooled to 4°C. The broken cells were pelleted at 2,500 g and washed with ice-cold Wash Buffer (20 mM Hepes-KOH pH 7.2 containing 0.15 M mannitol, 2 mM EDTA, 0.1% (w/v) BSA, and 1 mM MgCl_2_). After a 2 minutes 1000 g centrifugation, the pellet was resuspended in 2 ml of the same solution, stirred vigorously for 45 seconds, and then 6 ml of 20 mM Hepes-KOH buffer pH 7.2 containing 0.15 M mannitol, 0.8 mM EDTA, and 4 mM MgCl_2_ were added. Mitochondria were pelleted at 12,000 g for 10 minutes, resuspended in the same last buffer and then loaded on a discontinuous Percoll gradient (13 %/21 %/45 %) in MET buffer (280 mM Mannitol, 10 mM Tris-HCl pH 6.8, 0.5 mM EDTA, and 0.1 % BSA) and centrifuged for 60 minutes at 40,000 g. Purified mitochondria were recovered at the 45/21 interface, washed two times in MET buffer and flash frozen in liquid nitrogen and stored at -80°C.

Respiratory complexes were then purified as follows: purified mitochondria were re-suspended in Mitochondria Lysis Buffer (20 mM HEPES-KOH, pH 7.5, 75 mM KCl, 1 mM DTT, 5% Digitonin, supplemented with proteases inhibitors (C0mplete EDTA-free)) to a 5 mg/ml concentration, resuspended by pipetting and incubated for 30 min in 4°C. The lysate was then homogenized using a Dounce homogenizer. Lysate was clarified by centrifugation at 14,000 g, 10 min at 4°C. 1 mL of the lysed mitochondria was then loaded on a 10-40 % sucrose gradient in the same buffer (with 0.1% Digitonin) and run for 15 h at 31k rpm in a SW41 rotor. Fractions corresponding to respiratory complexes were automatically collected using a BioComp gradient fractionator, pelleted and re-suspended in buffer and analyzed by cryo-EM.

### Cryo-EM grid preparation

4 µl of full respiratory complexes at a concentration of 1 µg/µl of proteins (BSA equivalent measurement on Nanodrop) were applied onto a Quantifoil R2/1 300-mesh holey carbon grid coated with 2nm continuous carbon film. Grids were glow-discharged at 5W for 20 sec in a Gatan glow discharger. The sample was plunge frozen in liquid ethane with a Vitrobot Mark IV system (temperature = 4 °C, humidity 100%, blot force 5) with a wait time of 25 sec before blotting for 2.5 sec.

### Cryo-EM data collection

Data collection was performed using a 300kV Krios G4 electron microscope (Thermo Fisher Scientific) equipped with a cold field-emission gun, a Selectrix X energy filter and a prototype Falcon 4i detector with improved DQE. The energy filter was operated with a slit width of 10eV. Data were collected using aberration-free image shift (AFIS) as incorporated within the Thermo Fisher EPU software at a nominal defocus of 0.5 to 2 µm and at a magnification of 165,000x, yielding a calibrated pixel size of 0.737 Å/pixel at the specimen level. A total of 9,757 and 6,997 micrographs were recorded as EER files over the same session but on two different grids respectively, as a movie stack, with a total electron dose of 40 e^−^/Å².

### Single particle cryo-EM data processing

The entire processing pipeline was carried out in cryoSPARC (*87*) and the processing workflow is shown in fig. S4.

Pre-processing and particle picking were performed independently on the two sets of micrographs collected, and the resulting good particles were then merged. For dataset 1, after motion correction and CTF estimation 9,457 micrographs were kept, and 6,198 micrographs were kept for dataset 2. Particles were then picked using both a combination of blob picker, a template picker and Topaz, notably finding rare side views. Particles positions were extracted with a box size of 800 pixels, down sampled to 256 pixels (pixel size of 2.3031 Å) for faster processing and submitted to several rounds of 2D classifications, after which 141,939 particles remained. Particles were then further classified in 3D using the 3D Variability job and the best class, 83,443 particles, were selected and retracted with box size of 800 pixels, down sampled to 540 pixels (pixel size of 1.0919 Å). A global refinement reached 2.93 Å resolution. After Global and Local CTF refinement, resolution improved to 2.66 Å. Reference Based Motion Correction was then performed and further improved the resolution to 2.55 Å. Particles were then down sampled 588 pixels (pixel size of 1.0027) and another round of Global and Local CTF refinement at full resolution reached to 2.82 Å with a mask covering the entire supercomplex. Focused refinements were performed using a mask for the entire complex I (2.51 Å resolution), which was then further local refined on the matrix arm (2.34 Å resolution), the proximal membrane arm (2.30 Å resolution) and the distal membrane arm (2.21 Å resolution). The same was performed for complex III (2.46 Å resolution) and after recentering reached 2.39 Å. Initial focused refinement with a mask covering both complex IVs reached 3.01 Å resolution, and then 2.73 Å for the proximal complex IV (closest to the carbonic anhydrase domain of complex I) and 2.67 Å for the distal one. After recentering, they respectively reached 2.39 Å and 2.55 Å resolution.

### Model building and refinement

Focused refined cryo-EM maps (all at resolution better than 3 Å) for complex I, III and IV were used as inputs for ModelAngelo (*88*) *de novo* sequencing from the experimental data (without input sequence). Peptide chains obtained were then searched against the *Chlamydomonas reinhardtii* UNIPROT database (Taxon ID 3055) using BLAST (*89*) and the corresponding AlphaFold2 (*90*) models were retrieved for all the identified proteins. All proteins were identified, see Tables 2 and 3 for full composition. AlphaFold2 models were then matched to the ModelAngelo models using the Matchmaker tool in ChimeraX (*91*), and then further rigid-body fitted into their respective cryo-EM maps. The protein models were then manually inspected and refined in COOT (*92*). Ligands were then placed first by homology with PDB 8BPX and then lipids were manually placed, directly identified from the density. Water molecules were placed using the PHENIX phenix.douse tool and semi-automatically curated using the “Check Waters” tool in COOT. The different parts of the model were then automatically refined in PHENIX against the best resolved focus-refined maps using the phenix.real_space_refine tool and then again manually refined in COOT, iterated through several cycles (*93*). Chimeric maps were generated by using the Map Box option in PHENIX to cut out densities around the models (8 Å around the atoms). Maps were then fitted into the consensus maps in ChimeraX and combined using the ‘vop maximum’ command. For the model of the I_2_ III_4_ IV_6_ respirasome, the C2-expanded map was used to build a model of half a respirasome. To do so, models of CI, CIII and CIVs obtained from the SPA data were rigid-body fitted and adjusted when necessary. Then, two half models were fitted in the C2 map and further adjusted. Geometry of the models was validated using MolProbity (Table 1) (*94*).

### Figure preparation

Membrane segmentations for visualization were automatically performed using MemBrain-seg (*95*). All figures depicting segmentations and molecular models were prepared using ChimeraX (*96*), and tomogram views with map-backs of the molecular complexes were done using ArtiaX (*97*). Raw tomogram visualization was performed in IMOD (*79*).

## Authorcontributions

F.W. performed biochemical purifications, cryo-EM and cryo-ET data processing, model building and interpretation, and initial writing of the manuscript. R.K., X.Z. and A.K. performed cryo-FIB milling and collected cryo-ET data. A.K. collected single particle data. S.K. and R.D.R. preprocessed and reconstructed cryo-ET data. M.O. and S.K. helped troubleshoot the STA pipeline. F.W. and B.E. conceived the project, interpreted results, and finalized the manuscript.

## Acknowledgments

We thank Mohamed Chami and David Kalbermatter from the University of Basel BioEM facility for help operating the electron microscopes during the cryo-EM sample screening. We thank Thalia Salinas for advice on mitochondria purification, Irene Vercellino for advice on generating chimeric cryo-EM maps, and Asier González Seviné and Mihaela Zavolan for providing access to the BioComp gradient fractionator. We thank Wojciech Wietrzynski, Caitlyn McCafferty, and Sophie van Dorst for critical reading of the manuscript. F.W. was supported by the Swiss National Science Foundation with a Swiss Postdoc Fellowship (project 210561) and by the Alexander von Humboldt Foundation through the Humboldt Research Fellowship Programme for Postdocs. B.D.E. acknowledges funding from ERC consolidator grant “cryOcean” (fulfilled by the Swiss State Secretariat for Education, Research and Innovation, M822.00045) and bridging postdoctoral funds from the University of Basel Biozentum.

## Data Availability

The single particle cryo-EM maps of *C. reinhardtii* respirasome have been deposited at the Electron Microscopy Data Bank (EMDB) and models on the protein data bank (PDB). SPA of the I_1_III_2_IV_2_ partial respirasome: EMD-50202 and EMD-50249 (PDB 9F5X), complex I focused: EMD-50203 and EMD-50248 (PDB 9F5Y), complex I matrix focused: EMD-50207, complex I membrane proximal focused: EMD-50208, complex I membrane distal: EMD-50209, complex III focused: EMD-50204 (PDB 9F5Z), complex IV proximal focused: EMD-50205 (PDB 9F60), complex IV distal focused: EMD-50206 (PDB 9F61). *In situ* STA maps and models are also on the EMDB and PDB. STA of the native I_2_III_4_IV_6_ respirasome with C1 symmetry: EMD-50216, with C2 symmetry: EMD-50210 (PDB 9F62), respirasome C2 expanded: EMD-50211, respirasome C2 expanded focused complex I: EMD-50214, respirasome C2 expanded focused complex III: EMD-50215, mitoribosome: EMD-50213, putative prohibitin: EMD-50212. The raw cryo-ET data is available at the Electron Microscopy Public Image Archive (EMPIAR), accession code: EMPIAR-11830.

## CompetingInterests

The authors declare the following financial interests/personal relationships which may be considered as potential competing interests: M.O., S.K., X.Z., and A.K. are employees of Thermo Fisher Scientific. F.W., R.D.R., and B.D.E. declare no competing interests.

## References

1. M. W. Gray, Mitochondrial Evolution. Cold SpringHarbor Perspectivesin Biology 4, a011403 (2012).

2. P. Mitchell, Coupling of Phosphorylation to Electron and Hydrogen Transfer by a Chemi-Osmotic type of Mechanism. Nature 191, 144–148(1961).

3. J. S. Sousa, E. D’Imprima, J. Vonck, “Mitochondrial Respiratory Chain Complexes” (2018; http://link.springer.com/10.1007/978-981-10-7757-9_7), pp. 167–227.

4. A. Mühleip, S.E. McComas, A. Amunts, Structure of a mitochondrial ATPsynthasewith bound native cardiolipin. eLife 8 (2019).

5. A. Mühleip, R. Kock Flygaard, J. Ovciarikova, A. Lacombe, P. Fernandes, L. Sheiner, A. Amunts, ATPsynthase hexamer assemblies shape cristae of Toxoplasma mitochondria. Nat Commun 12, 120 (2021).

6. R. K. Flygaard, A. Mühleip, V. Tobiasson, A. Amunts, Type III ATP synthase is a symmetry-deviated dimer that induces membrane curvature through tetramerization. Nat Commun 11, 5342 (2020).

7. T. E. Spikes, M. G. Montgomery, J. E. Walker, Structure of the dimeric ATP synthase from bovine mitochondria. Proceedingsof the National Academy of Sciencesof the United States of America 117, 23519–23526 (2020).

8. K. Parey, C. Wirth, J. Vonck, V. Zickermann, Respiratory complex I — structure, mechanism and evolution. Current Opinion in Structural Biology 63, 1–9(2020).

9. L. Zhou, M. Maldonado, A. Padavannil, F. Guo, J. A. Letts, Structures of Tetrahymena’s respiratory chain reveal the diversity of eukaryotic core metabolism. Science 376, 831–839 (2022).

10. N. Klusch, J. Senkler, Ö. Yildiz, W. Kühlbrandt, H.-P. Braun, Aferredoxin bridge connects the two arms of plant mitochondrial complex I. ThePlant Cell 33, 2072–2091(2021).

11. T. Pánek, M. Eliáš, M. Vancová, J. Lukeš, H. Hashimi, Returning to the Fold for Lessons in Mitochondrial Crista Diversity and Evolution. Current Biology 30, R575–R588(2020).

12. I. Vercellino, L. A. Sazanov, SCAF1 drives the compositional diversity of mammalian respirasomes. Nat Struct Mol Biol, 1–11 (2024).

13. J. Gu, M. Wu, R. Guo, K. Yan, J. Lei, N. Gao, M. Yang, The architecture of the mammalian respirasome. Nature 537, 639–643(2016).

14. J. A. Letts, K. Fiedorczuk, L. A. Sazanov, The architecture of respiratory supercomplexes. Nature 537, 644–648 (2016).

15. M. Wu, J. Gu, R. Guo, Y. Huang, M. Yang, Structure of Mammalian Respiratory Supercomplex I 1 III 2 IV 1. Cell 167, 1598–1609.e10 (2016).

16. Z. He, M. Wu, H. Tian, L. Wang, Y. Hu, F. Han, J. Zhou, Y. Wang, L. Zhou, Euglena’s atypical respiratory chain adapts to the discoidal cristae and flexible metabolism. Nat Commun 15, 1628 (2024).

17. A. Mühleip, R. K. Flygaard, R. Baradaran, O. Haapanen, T. Gruhl, V. Tobiasson, A. Maréchal, V. Sharma, A. Amunts, Structural basis of mitochondrial membrane bending by the I–II–III2– IV2 supercomplex. Nature 615, 934–938(2023).

18. N. Klusch, M. Dreimann, J. Senkler, N. Rugen, W. Kühlbrandt, H.-P. Braun, Cryo-EMstructure of the respiratory I + III2supercomplex from Arabidopsisthaliana at 2 Åresolution. Nat Plants 9, 142–156 (2023).

19. M. Maldonado, F. Guo, J.A. Letts, Atomic structures of respiratory complex III2, complex IV, and supercomplex III2-IV from vascular plants. eLife 10, 1–34(2021).

20. M. Maldonado, Z. Fan, K. M. Abe, J. A. Letts, Plant-specific features of respiratory supercomplex I + III2 from Vigna radiata. Nat. Plants 9, 157–168(2023).

21. F. Wú, A. Mühleip, T. Gruhl, L. Sheiner, A. Maréchal, A. Amunts, Structure of the II2-III2-IV2 mitochondrial supercomplex from the parasite Perkinsusmarinus. bioRxiv[Preprint] (2024). 10.1101/2024.05.25.595893.

22. A. M. Hartley, N. Lukoyanova, Y. Zhang, A. Cabrera-Orefice, S. Arnold, B. Meunier, N. Pinotsis, A. Maréchal, Structure of yeast cytochrome c oxidase in a supercomplex with cytochrome bc1. Nat Struct Mol Biol 26, 78–83(2019).

23. W. Kühlbrandt, Structure and Mechanisms of F-Type ATP Synthases. Annual Review of Biochemistry 88, 515–549(2019).

24. H. Schägger, K. Pfeiffer, The Ratio of Oxidative Phosphorylation Complexes I–V in Bovine Heart Mitochondria and the Composition of Respiratory Chain Supercomplexes*. Journalof Biological Chemistry 276, 37861–37867(2001).

25. H. Eubel, J. Heinemeyer, H.-P. Braun, Identification and characterization of respirasomes in potato mitochondria. Plant physiology 134, 1450–9(2004).

26. H. Eubel, L. Jänsch, H.-P. Braun, New insights into the respiratory chain of plant mitochondria. Supercomplexes and a unique composition of complex II. Plant physiology 133, 274–86 (2003).

27. H. Schägger, K. Pfeiffer, Supercomplexes in the respiratory chains of yeast and mammalian mitochondria. EMBOJ 19, 1777–1783 (2000).

28. C. L. McCafferty, S. Klumpe, R. E. Amaro, W. Kukulski, L. Collinson, B. D. Engel, Integrating cellular electron microscopy with multimodal data to explorebiologyacross space and time. Cell 187, 563–584(2024).

29. E. Nogales, J. Mahamid, Bridgingstructural and cell biology with cryo-electron microscopy. Nature 628, 47–56 (2024).

30. D. Tamborrini, Z. Wang, T. Wagner, S. Tacke, M. Stabrin, M. Grange, A. L. Kho, M. Rees, P. Bennett, M. Gautel, S. Raunser, Structure of the native myosin filament in the relaxedcardiac sarcomere. Nature 623, 863–871(2023).

31. H. Xing, R. Taniguchi, I. Khusainov, J.P. Kreysing, S. Welsch, B. Turoňová, M. Beck, Translation dynamics in human cells visualizedat highresolution revealcancer drugaction. Scienc*e***381**, 70–75 (2023).

32. L. Xue, S. Lenz, M. Zimmermann-Kogadeeva, D. Tegunov, P. Cramer, P. Bork, J. Rappsilber, J. Mahamid, Visualizing translation dynamics at atomic detail inside a bacterial cell. Nature 610, 205–211 (2022).

33. M. Schaffer, B. Engel, T. Laugks, J. Mahamid, J. Plitzko, W. Baumeister, Cryo-focused Ion Beam Sample Preparation for Imaging Vitreous Cells by Cryo-electron Tomography. BIO-PROTOCOL5 (2015).

34. L. N. Young, E. Villa, BringingStructure to Cell Biologywith Cryo-Electron Tomography. Annu. Rev. Biophys. 52, 573–595 (2023).

35. S. Dupuis, S. S. Merchant, Chlamydomonas reinhardtii: a model for photosynthesis and so much more. Nature Methods 20, 1441–1442(2023).

36. H. van den Hoek, N. Klena, M. A. Jordan, G. Alvarez Viar, R. D. Righetto, M. Schaffer, P. S. Erdmann, W. Wan, S. Geimer, J.M. Plitzko, W. Baumeister, G. Pigino, V. Hamel, P. Guichard, B. D. Engel, In situ architecture of the ciliary base reveals the stepwise assembly of intraflagellar transport trains. *Science (New York*, N.Y.) 377, 543–548(2022).

37. M. Gui, X. Wang, S.K. Dutcher, A. Brown, R. Zhang, Ciliary central apparatus structure reveals mechanisms of microtubule patterning. Nat Struct Mol Biol 29, 483–492(2022).

38. F. Waltz, T. Salinas-Giegé, R. Englmeier, H. Meichel, H. Soufari, L. Kuhn, S. Pfeffer, F. Förster, B. D. Engel, P. Giegé, L. Drouard, Y. Hashem, How to build a ribosome from RNAfragments in Chlamydomonas mitochondria. Nature Communications 12, 7176 (2021).

39. P. Cardol, D. González-Halphen, A. Reyes-Prieto, D. Baurain, R.F. Matagne, C. Remacle, The Mitochondrial Oxidative Phosphorylation Proteome of Chlamydomonas reinhardtii Deduced from the Genome Sequencing Project. Plant Physiol 137, 447–459(2005).

40. J. Findinier, L.-M. Joubert, M. F. Schmid, A. Malkovskiy, W. Chiu, A. Burlacot, A. R. Grossman, Dramatic Changes in Mitochondrial Subcellular Location and Morphology Accompany Activation of the CO2 Concentrating Mechanism. bioRxiv [Preprint] (2024). 10.1101/2024.03.25.586705.

41. P. Ježek, M. Jabůrek, B. Holendová, H. Engstová, A. Dlasková, Mitochondrial Cristae Morphology Reflecting Metabolism, Superoxide Formation, Redox Homeostasis, and Pathology. Antioxid Redox Signal 39, 635–683(2023).

42. S. E. Siegmund, R. Grassucci, S. D. Carter, E. Barca, Z. J. Farino, M. Juanola-Falgarona, P. Zhang, K. Tanji, M. Hirano, E. A. Schon, J. Frank, Z. Freyberg, Three-Dimensional Analysis of Mitochondrial Crista Ultrastructure in a Patient with Leigh Syndromeby In Situ Cryoelectron Tomography. iScience 6, 83–91(2018).

43. B. A. Barad, M. Medina, D. Fuentes, R. L. Wiseman, D. A. Grotjahn, Quantifying organellar ultrastructure in cryo-electron tomographyusing a surface morphometrics pipeline. Journal of Cell Biology 222, e202204093 (2023).

44. F. Lange, M. Ratz, J.-N. Dohrke, D. Wenzel, P. Ilgen, D. Riedel, S. Jakobs, In-situ architecture of the human prohibitin complex. bioRxiv [Preprint] (2024). 10.1101/2024.02.14.579514.

45. L. Dietrich, A.-N. A. Agip, A. Schwarz, C. Kunz, W. Kühlbrandt, In situ structure and rotary states of mitochondrial ATP synthase in whole cells. bioRxiv [Preprint] (2024). 10.1101/2024.03.27.586927.

46. E. Buzzard, M. McLaren, P. Bragoszewski, A. Brancaccio, H. C. Ford, B. Daum, P. Kuwabara, I. Collinson, V. A. M. Gold, Theconsequence of ATPsynthase dimer angle on mitochondrial morphology studied by cryo-electron tomography. Biochemical Journal 481, 161–175(2024).

47. K. M. Davies, C. Anselmi, I. Wittig, J.D. Faraldo-Gómez, W. Kühlbrandt, Structure of the yeast F1Fo-ATPsynthase dimer and its role in shaping the mitochondrial cristae. Proceedings of the National Academy of Sciences 109, 13602–13607(2012).

48. S. Pfeffer, M. W. Woellhaf, J. M. Herrmann, F. Förster, Organization of the mitochondrial translation machinery studied in situ by cryoelectron tomography. Nature Communications 6, 60199 (2015).

49. R. Englmeier, S. Pfeffer, F. Förster, Structure of the Human Mitochondrial Ribosome Studied In Situ by Cryoelectron Tomography. Structure 25, 1574–1581.e2 (2017).

50. N. Pfanner, M. Meĳer, Mitochondrial biogenesis: TheTom and Tim machine. Current Biology 7, R100–R103(1997).

51. W. Zheng, P. Chai, J. Zhu, K. Zhang, High-resolution in situ structures of mammalian respiratory supercomplexes. Nature, 1–8 (2024).

52. K. M. Davies, T. B. Blum, W. Kühlbrandt, Conservedin situ arrangement of complex I and III2 in mitochondrial respiratory chain supercomplexes of mammals, yeast, and plants. Proceedingsof the National Academyof Sciencesof the United Statesof America 115, 3024– 3029 (2018).

53. H.-P. Braun, N. Klusch, Promotion of oxidative phosphorylation by complex I-anchored carbonic anhydrases? Trends in Plant Science 29, 64–71(2024).

54. H. Soufari, C. Parrot, L. Kuhn, F. Waltz, Y. Hashem, Specific features and assembly of the plant mitochondrial complex I revealed by cryo-EM. Nature Communications 11, 5195 (2020).

55. J.A. Letts, K. Fiedorczuk, G. Degliesposti, M. Skehel, L. A. Sazanov, Structures of Respiratory Supercomplex I+III2 Reveal Functional and Conformational Crosstalk. Molecular Cell 75, 1131–1146.e6 (2019).

56. S. Shimada, K. Shinzawa-Itoh, J. Baba, S. Aoe, A. Shimada, E. Yamashita, J. Kang, M. Tateno, S. Yoshikawa, T. Tsukihara, Complex structure of cytochrome c–cytochrome c oxidase reveals a novel protein–protein interaction mode. The EMBOJournal 36, 291–300(2017).

57. S. R. N. Solmaz, C. Hunte, Structure of Complex III with Bound Cytochrome c in Reduced State and Definition of a Minimal Core Interface for Electron Transfer*. Journalof Biological Chemistry 283, 17542–17549(2008).

58. A. Kohler, A. Barrientos, F. Fontanesi, M. Ott, The functional significance of mitochondrial respiratory chain supercomplexes. EMBOreports 24, e57092 (2023).

59. I. Vercellino, L. A. Sazanov, The assembly, regulation and function of the mitochondrial respiratory chain. Nature ReviewsMolecular Cell Biology, 1–21(2021).

60. J. A. Letts, L. A. Sazanov, Clarifying the supercomplex: the higher-order organization of the mitochondrial electron transport chain. Nature Structural & Molecular Biology 24, 800–808 (2017).

61. J. S. Sousa, D. J. Mills, J. Vonck, W. Kühlbrandt, Functional asymmetry and electron flow in the bovine respirasome. eLife 5 (2016).

62. F. Diaz, H. Fukui, S. Garcia, C. T. Moraes, Cytochrome c oxidase is required for the assembly/stability of respiratory complex I in mouse fibroblasts. Mol Cell Biol 26, 4872–4881 (2006).

63. E. Lapuente-Brun, R. Moreno-Loshuertos, R. Acín-Pérez, A. Latorre-Pellicer, C. Colás, E. Balsa, E. Perales-Clemente, P. M. Quirós, E. Calvo, M. A. Rodríguez-Hernández, P. Navas, R. Cruz, Á. Carracedo, C. López-Otín, A. Pérez-Martos, P. Fernández-Silva, E. Fernández-Vizarra, J. A. Enríquez, Supercomplex Assembly Determines Electron Flux in the Mitochondrial Electron Transport Chain. Science 340, 1567–1570(2013).

64. J. G. Fedor, J. Hirst, Mitochondrial Supercomplexes Do Not Enhance Catalysis by Quinone Channeling. Cell Metabolism 28, 525–531.e4 (2018).

65. Y. Chaban, E. J. Boekema, N. V. Dudkina, Structures of mitochondrial oxidative phosphorylation supercomplexes and mechanisms for their stabilisation. Biochimica et Biophysica Acta (BBA)-Bioenergetics 1837, 418–426 (2014).

66. M. C. Pöverlein, A. Jussupow, H. Kim, V. R. I. Kaila, Protein-Induced Membrane Strain Drives Supercomplex Formation. bioRxiv [Preprint] (2024). 10.1101/2024.07.13.602826.

67. D. Milenkovic, J. Misic, J. F. Hevler, T. Molinié, I. Chung, I. Atanassov, X. Li, R. Filograna, A. Mesaros, A. Mourier, A. J. R. Heck, J. Hirst, N.-G. Larsson, Preserved respiratory chain capacity and physiology in mice with profoundly reduced levels of mitochondrial respirasomes. Cell Metabolism 35, 1799–1813.e7 (2023).

68. M. Brischigliaro, A. Cabrera-Orefice, S. Arnold, C. Viscomi, M. Zeviani, E. Fernández-Vizarra, Structural rather than catalytic role for mitochondrial respiratory chain supercomplexes. eLife 12 (2023).

69. K. M. Davies, M. Strauss, B. Daum, J.H. Kief, H. D. Osiewacz, A. Rycovska,V. Zickermann, W. Kuhlbrandt, Macromolecular organization of ATP synthase and complex I in whole mitochondria. Proceedingsof the National Academyof Sciences 108, 14121–14126(2011).

70. W. Wietrzynski, M. Schaffer, D. Tegunov, S. Albert, A. Kanazawa, J.M. Plitzko, W. Baumeister, B. D. Engel, Charting the native architecture of chlamydomonas thylakoid membranes with single-molecule precision. eLife 9 (2020).

71. J. Gu, T. Liu, R. Guo, L. Zhang, M. Yang, The coupling mechanism of mammalian mitochondrial complex I. Nat Struct Mol Biol 29, 172–182(2022).

72. V. Kravchuk, O. Petrova, D. Kampjut, A. Wojciechowska-Bason, Z. Breese, L. Sazanov, A universal coupling mechanism of respiratory complex I. Nature 609, 808–814(2022).

73. D. Kampjut, L. A. Sazanov, The coupling mechanism of mammalian respiratory complex I. Science 370, eabc4209 (2020).

74. K. Parey, J. Lasham, D. J. Mills, A. Djurabekova, O. Haapanen, E. G. Yoga, H. Xie, W. Kühlbrandt, V. Sharma, J. Vonck, V. Zickermann, High-resolution structure and dynamics of mitochondrial complex I—Insightsinto the proton pumping mechanism. Science Advances 7, eabj3221 (2021).

75. S. Khavnekar, P. S. Erdmann, W. Wan, TOMOMAN:a software package for large scale cryo-electron tomography data preprocessing, community data sharing, and collaborative computing. bioRxiv[Preprint] (2024).10.1101/2024.05.02.589639.

76. S. Q. Zheng, E. Palovcak, J.-P. Armache, K. A. Verba, Y. Cheng, D. A. Agard, MotionCor2: anisotropic correction of beam-induced motion for improved cryo-electron microscopy. Nature Methods 14, 331–332(2017).

77. A. Rohou, N. Grigorieff, CTFFIND4:Fast and accurate defocus estimation from electron micrographs. Journalof Structural Biology 192, 216–221(2015).

78. S. Zheng, G. Wolff, G. Greenan, Z. Chen, F. G. A. Faas, M. Bárcena, A. J. Koster, Y. Cheng, D. A. Agard, AreTomo: An integrated software package for automated marker-free, motion-corrected cryo-electron tomographic alignment and reconstruction. Journal of Structural Biology: X 6, 100068 (2022).

79. Computer Visualization of Three-Dimensional Image Data Using IMOD. https://bio3d.colorado.edu/imod/paper/.

80. B. Turoňová, F. K. M. Schur, W. Wan, J. A. G. Briggs, Efficient 3D-CTFcorrection for cryo-electron tomography using NovaCTFimproves subtomogram averagingresolution to 3.4 Å. Journalof Structural Biology 199, 187–195(2017).

81. T. O. Buchholz, M. Jordan, G. Pigino, F. Jug,Cryo-CARE:Content-Aware ImageRestoration for Cryo-Transmission Electron Microscopy Data. Proceedings - International Symposium on Biomedical Imaging 2019-April, 502–506(2018).

82. W. Wan, S. Khavnekar, J. Wagner, STOPGAP: anopen-source packagefor template matching, subtomogram alignment and classification. Acta Cryst D 80 (2024).

83. T. Hrabe, Y. Chen, S. Pfeffer, L. Kuhn Cuellar, A.-V. Mangold, F. Förster, PyTom: A python-based toolbox for localization of macromolecules in cryo-electron tomograms and subtomogram analysis. Journalof Structural Biology 178, 177–188(2012).

84. M. L. Chaillet, G. van der Schot, I. Gubins, S. Roet, R. C. Veltkamp, F. Förster, Extensive Angular Sampling Enables the Sensitive Localization of Macromolecules in Electron Tomograms. International Journalof Molecular Sciences 24, 13375 (2023).

85. J. Zivanov, J. Otón, Z. Ke, A. von Kügelgen, E. Pyle, K. Qu, D. Morado, D. Castaño-Díez, G. Zanetti, T. A. Bharat, J. A. Briggs, S. H. Scheres, A Bayesian approach to single-particle electron cryo-tomography in RELION-4.0.eLife 11, e83724 (2022).

86. S. L. Ilca, A. Kotecha, X. Sun, M. M. Poranen, D. I. Stuart, J. T. Huiskonen, Localized reconstruction of subunits from electron cryomicroscopy images of macromolecular complexes. Nat Commun 6, 8843 (2015).

87. A. Punjani, J. L. Rubinstein, D. J. Fleet, M. A. Brubaker, cryoSPARC: algorithms for rapid unsupervised cryo-EMstructure determination. Nature Methods 14, 290–296(2017).

88. K. Jamali, L. Käll, R. Zhang, A. Brown, D. Kimanius, S. H. W. Scheres, Automated model building and protein identification in cryo-EMmaps. Nature, 1–8(2024).

89. S. F. Altschul, T. L. Madden, A. A. Schäffer, J. Zhang, Z. Zhang, W. Miller, D. J. Lipman, Gapped BLASTandPSI-BLAST:anew generation of protein database search programs. Nucleic acids research 25, 3389–402(1997).

90. J. Jumper, R. Evans, A. Pritzel, T. Green, M. Figurnov, O. Ronneberger, K. Tunyasuvunakool, R. Bates, A. Žídek, A. Potapenko, A. Bridgland, C. Meyer, S. A. A. Kohl, A. J. Ballard, A. Cowie, B. Romera-Paredes, S. Nikolov, R. Jain, J. Adler, T. Back, S. Petersen, D. Reiman, E. Clancy, M. Zielinski, M. Steinegger, M. Pacholska, T. Berghammer, S. Bodenstein, D. Silver, O. Vinyals, A. W. Senior, K. Kavukcuoglu, P. Kohli, D. Hassabis, Highly accurate protein structure prediction with AlphaFold. Nature 596, 583–589(2021).

91. E. C. Meng, T. D. Goddard, E. F. Pettersen, G. S. Couch, Z. J. Pearson, J.H. Morris, T. E. Ferrin, UCSFChimeraX: Tools for structure building and analysis. Protein Science 32, e4792 (2023).

92. P. Emsley, B. Lohkamp, W. G. Scott, K. Cowtan, Features and development of Coot. Acta Crystallographica Section DBiological Crystallography 66, 486–501(2010).

93. D. Liebschner, P. V. Afonine, M. L. Baker, G. Bunkóczi, V. B. Chen, T. I. Croll, B. Hintze, L.-W. Hung, S. Jain, A. J. McCoy, N. W. Moriarty, R. D. Oeffner, B. K. Poon, M. G. Prisant, R. J. Read, J. S. Richardson, D. C. Richardson, M. D. Sammito, O. V. Sobolev, D. H. Stockwell, T. C. Terwilliger, A. G. Urzhumtsev, L. L. Videau, C. J. Williams, P. D. Adams, Macromolecular structure determination using X-rays, neutrons and electrons: recent developments in Phenix. Acta Crystallographica Section DStructural Biology 75, 861–877(2019).

94. C. J. Williams, J.J. Headd, N. W. Moriarty, M. G. Prisant, L. L. Videau, L. N. Deis, V. Verma, D. A. Keedy, B. J.Hintze, V. B. Chen, S. Jain,S. M. Lewis, W. B.Arendall, J.Snoeyink, P. D. Adams, S. C. Lovell, J. S. Richardson, D. C. Richardson, MolProbity: More and better reference data for improved all-atom structure validation. Protein Sci 27, 293–315(2018).

95. L. Lamm, S. Zufferey, R. D. Righetto, W. Wietrzynski, K. A. Yamauchi, A. Burt, Y. Liu, H. Zhang, A. Martinez-Sanchez, S. Ziegler, F. Isensee, J.A. Schnabel, B. D. Engel, T. Peng, MemBrain v2: an end-to-end tool for the analysis of membranes in cryo-electron tomography. bioRxiv [Preprint] (2024).10.1101/2024.01.05.574336.

96. T. D. Goddard, C. C. Huang, E. C. Meng, E. F. Pettersen, G. S. Couch, J.H. Morris, T. E.Ferrin, UCSFChimeraX: Meeting modern challenges in visualization and analysis. Protein Science 27, 14–25 (2018).

97. U. H. Ermel, S. M. Arghittu, A. S. Frangakis, ArtiaX: An electron tomography toolbox for the interactive handling of sub-tomograms in UCSFChimeraX. Protein Sci 31, e4472 (2022).

